# Multi-region spatial transcriptome analysis reveals cellular networks and pathways associated with hepatocellular carcinoma recurrence

**DOI:** 10.1101/2023.07.26.549242

**Authors:** Aziz Aiderus, Pratap Veerabrahma Seshachalam, Khaireen Idzham, Matias Caldez, Raghuvaran Shanmugam, Ita Novita Sari, Dorcas Hei Hui Ying, Shay Lee Chong, Karthik Sekar, Sin Chi Chew, Gao Bin Chen, Alexander Yaw-Fui Chung, Peng Chung Cheow, Juinn Huar Kam, Alfred Wei-Chieh Kow, Iyer Shridhar Ganpathi, Shihleone Loong, Wei-Qiang Leow, Kaina Chen, Rawisak Chanwat, Vanessa H. de Villa, Peng Soon Koh, Glenn K Bonney, Brian K. P. Goh, Wai Leong Tam, Vinay Tergaonkar, Pierce Kah Hoe Chow

**Author notes:** **Corresponding author**: Pierce Kah Hoe Chow. These authors contributed equally to this work (Aziz Aiderus, Pratap Veerabrahma Seshachalam).

## Abstract

Hepatocellular carcinomas (HCC) are driven by various etiologies and molecular diversity at presentation. Patient prognosis post-surgery is generally dismal, and the majority respond poorly to adjuvant targeted and/or immuno-therapies. Tumours are an ecosystem comprised of organization and interaction between different cell types that may contribute to clinically significant outcomes, such as disease recurrence. To better understand this phenomenon, we leveraged on a local cohort of patients with or without recurrence to generate spatial transcriptome profiles from multiple sectors from each tumour. We identified widespread gene expression intra- and inter tumour heterogeneity observed across the tumour sectors. Our analysis also revealed the cell type enrichment and localization, and ligand-receptor interactions identify a specific subset of endothelial cell enriched in primary tumours of patients with recurrence. Altogether, this study describes the spatial gene expression landscape in HCC patients associated with disease recurrence.

## Introduction

Liver cancer is the sixth most diagnosed cancer and third most common cause of worldwide cancer deaths [1]. Approximately 85% of liver cancers are hepatocellular carcinoma and only approximately 20% of HCC is diagnosed at an early stage where potentially curative therapies (surgical resection, liver transplantation, radiofrequency ablation) are feasible [2]. This malignancy develops from multiple etiological factors, including chronic hepatitis B/C infection and metabolic dysfunction (e.g., non-alcoholic fatty liver disease) [3]. Surgical resection for patients with HCC is hampered by a high frequency of disease recurrence (between 50-70%) that occurs in a bi-modal pattern with an early peak at 1 year primarily from micro-metastases from the index tumour and a second peak between 4-5 years with de-novo cancer arising from the drivers within the underlying liver disease [4, 5]. As a result, surgical resection only confers a modest 5-year overall survival (OS) of > 60% and a poor recurrence-free survival of 35%, thereby presenting an urgent need to clarify factors driving recurrence [6–8]. Efficacious adjuvant therapy after curative intent surgical resection in HCC has just become available recently (https://doi.org/10.1158/1538-7445.AM2023-CT003) but there are still no predictive biomarkers of HCC and a granular understanding of changes in the tumour micro-environment that predicts early recurrence remains unclear [9]. Single cell RNA sequencing has been employed in several HCC studies to elucidate the cell types and states present within the tumour ecosystem and how it may be associated with clinically significant features, such recurrence and treatment response [10–17]. For example, Sun *et al* performed single cell RNA sequencing analysis comparing HCC tumours from 12 treatment-naïve and 6 early-relapse patients and observed that recurrent tumours exhibited enrichment of CD8+ T cells with low cytotoxicity which may result in poor anti-tumour immunity and disease relapse [16]. However, tumours are an ecosystem that comprise multiple cell types that interact with one another in diverse environments, such as altered vasculature, immune predation, and nutrient status [18]. Furthermore, the organisation of these cell compartments may influence cell-cell interactions and tumour biology. For instance, the presence of tertiary lymphoid structures (TLS), which are localised aggregates of immune cells within the tumour, is associated with response to immune checkpoint blockade in several cancer types [19]. Therefore, a cellular and molecular map at spatial resolution of HCC tumours with features of clinical interest, such as recurrence, may complement our current understanding of disease biology.

The 10X Genomics Visium platform provides an unbiased spatial transcriptome platform that uses immobilised poly (dT) probes on a glass slide with spatial barcodes to hybridise mRNA released from a tissue section after permeabilization. Each slide consists of 5,000 spots, each approximately 55 µm in diameter, which can accommodate several cells. To date, two studies have employed the Visium platform describe the spatial architecture of primary liver cancers, including hepatocellular carcinoma and intrahepatic cholangiocarcinoma [20, 21]. These studies provide a clear indication of the high levels of transcriptomic and copy number variation observed across each tumour section. However, single sampling of tumours has been shown to underappreciate tumour heterogeneity, and analysis of spatially distinct sites further unravelled the degree of genomic heterogeneity present within a tumour [22], which may have implications for inferring tumour characteristics, treatment response, and prognosis [23]. In particular, how the spatial organisation of the tumour ecosystem influences patient outcome after surgical resections remains to be determined and may provide actionable clinical insights.

The Precision Medicine in Liver Cancer across an Asia-Pacific Network (PLANet 1.0, NCT03267641) was conceptualised in 2016 to address the urgent unmet need for predictive biomarkers in HCC. This multinational, multi-disciplinary, longitudinal study was based on a strategy of multi-region sampling of resected HCC, and subsequent biopsy of recurrent tumours to perform comprehensive genomics and immunomics analysis. This study is among the leading prospective cohort worldwide, where the well annotated clinical and associated multi-omics data describe the intratumoral heterogeneous landscape and the natural trajectory of disease [24, 25] (https://doi.org/10.1016/j.jhepr.2023.100715). Patients were followed closely after surgery until recurrence or death, or up to three years after surgical resection. In this study, we leveraged on patient material from the PLANet 1.0 study to perform unbiased spatial transcriptomic analysis. This study is the first to perform spatial transcriptomic analysis on multiple distinct regions of a tumour, which allowed us to explore the tumour ecosystem heterogeneity and describe the cell types, interaction networks, ligand-receptor interactions, and how candidate biomarkers identified in this study correlates with survival. These findings provide a useful spatial cellular and molecular atlas to better appreciate the natural course of HCC post-surgery.

## Experimental procedures

### Patient samples for 10X Genomics Visium spatial transcriptomics

Patients with primary HCC underwent surgical resection at National Cancer Centre Singapore (N=1), Singapore General Hospital (N=1) and National University Hospital Singapore (N=2) were recruited. These four patients were a subset of the PLANet study (NCT03267641) conducted under the auspices of the Asia-Pacific Hepatocellular Carcinoma (AHCC) trials group. PLANET recruitment criteria required the patients to have no extra-hepatic metastasis (defined as lymph node <2 cm, lung modules < 1 cm, farther lymph nodes < 2 cm) with R0 or R1 resection and Child-Pugh ≤ 7 points without clinical ascites. A full set of patient recruitment criteria can be found in the Supplementary Notes from our previous report [24]. This study was approved by the Central Institution Review Board of SingHealth (2016/2626 and 2018/2112) and written consent from patients were obtained.

### Tumour sampling methodology

The resected specimens were acquired immediately and transported on ice in a temperature controlled cooler container to a pathologist, where each specimen was subjected to multi-region sampling [24]. Non-tumour normal tissues were defined as segments at least 2 cm away from the tumour. The tumour was then cut through the capsule, photographed, and inspected for fibrosis, necrosis, haemorrhage and cystic changes. Then, multiple sectors (regions) along one axis of a single tumor slice were then harvested. After surgery and pathology evaluation, samples were delivered in MACS buffer on ice. Upon arrival at the lab, samples were removed from buffer, cut to approximately 4-5 mm^3^ pieces and embedded in optimal cutting temperature compound.

### Collection and preparation of primary tumour samples

Surgically resected samples were placed in MACS buffer and transported to the laboratory. At arrival samples were immediately embedded in optimum cutting temperature (OCT) compound in a mold and frozen on dry ice, before storing in −80°C. Prior to performing tissue optimisation, 10 consecutive sections from each sample were obtained, RNA extracted, and RNA Integrity Number analysis performed using the Bioanalyzer RNA Nano or Pico Chip. Only samples with RIN of ≥ 7.0 were taken further for spatial transcriptomic analysis on the 10X Genomics Visium platform.

### Tissue fixation, H&E staining, and imaging

Once sectioned onto the Visium slide, tissue sections were fixed in methanol for 30 mins at −20°C. The slide was then immersed in isopropanol for 1 min, and hematoxylin for 7 mins, Bluing buffer for 2 mins and Eosin mix for 1 min at room temperature. The slides were washed in ultrapure water after each staining step and finally, were incubated at 37°C on a heat block for one minute to dry. Coverslips were mounted using 85% glycerol and then imaged on Zeiss microscrope at 20X magnification, as recommended by 10X Genomics.

### Determining optimal tissue permeabilization conditions to ensure maximal mRNA recovery

For each tissue, the permeabilisation time was determined to ensure maximum recovery of mRNA. Tissue sections were placed on the Tissue Optimization slide and fixed and stained with H&E reagents prior to imaging. Before tissue permeabilization, tissues were treated with 75 µL of Liberase^TM^ for 15 mins at 37°C. The enzyme mix was removed, and tissues washed with 100 µL 0.1X sodium saline citrate (SSC) buffer before proceeding with tissue permeabilization. Seventy µL of permeabilization enzyme was added to each tissue for 5-, 10-, and 15 minutes. Permeabilization results in the release of mRNA onto a poly(T) oligonucleotide on the capture areas and cDNA was then synthesized and the fluorescent signal determined. High quality RNA from the HEK293T cell line were used as positive control. The efficiency of permeabilization was determined as the time required to achieve the strongest fluorescent intensity observed with minimal background. After cDNA synthesis, the tissue slide was immersed in 0.2X SSC buffer with 0.1% SDS, followed by 0.2X SSC buffer, and finally 0.1X SSC buffer. Fluorescence was imaged on the Cy3 channel at 10X magnification, and the time point with the most even and brightest fluorescence was determined at the optimal permeabilization condition. Based on this time point, multi-sector sectioning was performed at a thickness of 10-14 µm onto a Visium slide and processed further.

### Tissue permeabilization, reverse transcription, second strand synthesis and denaturation, and cDNA amplification

The stained slide was then placed in the Slide Cassette and permeabilization enzyme added for the optimised time point. The poly(A) mRNA released were captured by poly(dT) primers on the spots and washed with SSC buffer. Reverse transcription was then performed using the master mix provided containing the reagents on the Thermocycler Adaptor provided to generate spatially barcoded full-length cDNA from polyA mRNA recovered. The cDNA was removed from the slide by incubating in 0.08 M potassium hydroxide for 5 mins and samples from each well transferred to a strip-tube with Tris-HCl (1M, pH 7.0). Next, 1 µL of the denatured sample was utilized for qPCR amplification to determine the sample C_q_ value. Next, cDNA synthesis was performed with the denatured sample using the cDNA Amplification Mix provided and this mix was placed in the Thermocycler Adaptor for one cycle. The resulting cDNA was cleaned up by according to instructions provided by 10X Genomics, involving incubation with SPRIselect beads and ethanol washes, before eluting in EB buffer. The quantity and quality of the cDNA was determined using Agilent Bioanalyzer High Sensitivity Chip, and the cDNA was diluted tenfold prior to loading onto the chip.

### Illumina sequencing library generation

Sequencing library was prepared according to the protocol provided by 10X Genomics. Briefly, using 10 µL of cDNA from the previous step was taken further for enzymatic fragmentation, end-repair and A-tailing, and size selection performed using SPRIselect beads. Sequencing adaptors were ligated, double-sided size selection and clean up performed, and sample index (Dual Index Kit TT Set A) PCR was conducted before a final double-sided size selection and clean up. Library quality and quantity was assessed using the Agilent High Sensitivity DNA Chip and final quantitation performed using Qubit. Sequencing libraries were then submitted to a local sequencing partner and sequenced at a depth of 25,000 read pairs per spot multiplied by the estimated fudicial frame coverage of the tissue on the Illumina HiSeq platform.

### Tumour clusters and survival analysis

To determine whether spatial gene expression clusters were associated with disease survival, we generated a bulk gene expression profile and applied them to bulk RNA-Seq data from 91 patients in the PLANet cohort (including the four patients in this study) for survival analysis. These patients have either developed HCC recurrence, have died or been followed-up for at least three years without recurrence. The average gene expression values from each cluster were generated and we focused on genes that were previously shown to be significantly associated with overall survival.

### Raw data preprocessing

Visium spatial RNA sequenced reads were analysed using the space ranger pipeline (version: 1.3.0, 10X genomics) to align the reads to hg38 reference genome and generate gene-spot matrixes corresponding to H & E image from each tissue. The Visium has combined spatial algorithms with widely used RNA-seq aligner STAR. Initial inspection of data was done using Loupe browser (6.3.0, 10X Genomics).

### Quality control

The filtered featured matrix (UMI counts), spatial coordinates csv file, scalefactor json file and H& E image were imported into python package squidpy using sc.read_visium function. Quality control was performed to remove low quality spots that consists of less than 200 genes or mitochondrial percentage > 5 %. Genes were removed which were expressed in less than 3 spots. Mitochondrial and ribosomal genes were excluded.

### Normalisation, dimension reduction, clustering, and ligand-receptor analysis

Gene expression count data is normalized initially using sc.pp.normalize_total and subsequently followed by log1p normalization from squidpy package. The top 2000 highly variable genes were selected based on expression mean and variance. Normalised gene expression data was subjected to principal component analysis (PCA) using the first 30 PCA components, and shared nearest neighborhood graphing and Leiden clustering performed. Leiden algorithm is improved version of Louvain algorithm and outperforms other clustering algorithms for single cell data sets [26, 27]([Du et al., 2018]). Therefore, we used Leiden clustering to cluster the spots based on gene expression. Dimensionality reduction was done using Uniform Manifold Approximation and Projection (UMAP) and squidpy implemented for clustering and dimension reduction analysis. Neighbourhood enrichment analysis of clusters was performed by squidpy function sq.gr.spatial_neighbours and differential gene expression analysis was performed via Wilcoxon rank sum statistical test on each cluster. Ligand – receptor analysis was performed using the squidpy package and the omnipath database was nominated for ligand – receptor annotation. In simple terms, it is a permutation test for ligand-receptor expression across different spatial gene expression clusters.

## Results

### Experimental design, clinico-pathological characteristics, and clinical trajectory of the study cohort

To determine the spatial molecular and cellular ecosystem of HCC, we conducted spatial transcriptome analysis on multiple sectors per tumour from patients with (n = 3) or without recurrence (n = 1), with well annotated clinical data (**Fig 1A**). This cohort was part of a previous PLANet study [24] and include both hepatitis B-positive and negative individuals. Where possible, as reference, we profiled adjacent non-tumour tissue and defined them as ‘normal’. Each tumour was sampled at 2-5 distinct regions, and at least one section per region was obtained for spatial transcriptomic analysis.

**Figure 1.**
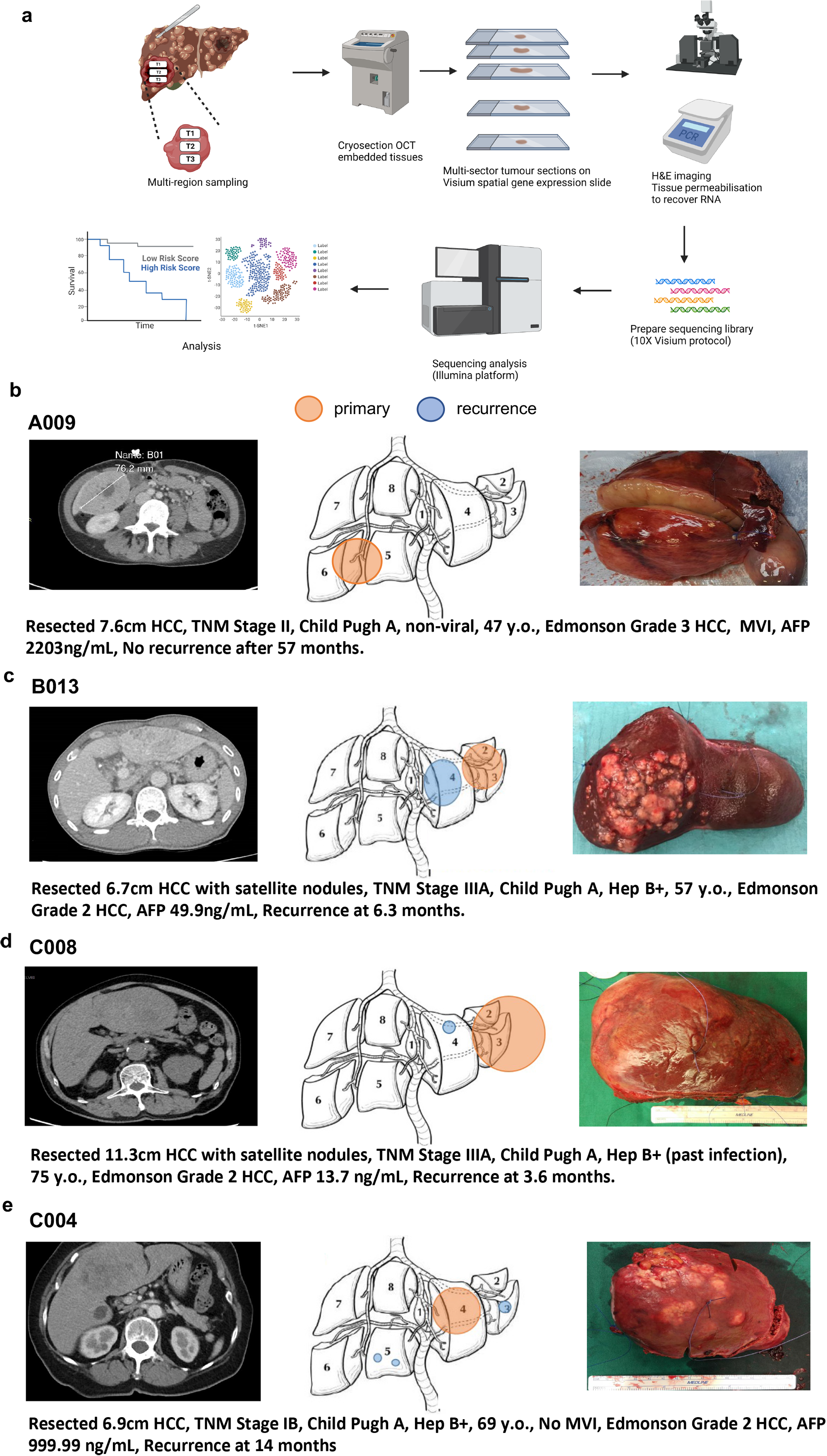
Study design and patient cohort. (a) Schematic of processing tissues for the 10X Genomics Visium spatial transcriptomic analysis. Sectors marked and obtained from a tumour tissue that was grided during pathology evaluation. Tissues were snapped frozen and embedded in OCT compound. Fourteen-micron sections were performed onto a tissue optimisation slide to determine the permeabilization time to maximise RNA yield. Tissue sections were then placed on the gene expression slide and RNA recovered, before generating libraries for sequencing and downstream bioinformatic analysis. (b-e) Computed tomography, schema of tumour location, and gross tumour images of patients A009, B013, C008, C004.

The 4 patients from whom biosamples were collected from their surgically resected HCC formed the clinical cohort. All 4 patients were Child-Pugh A, received R0 resection (resection margins were free from macroscopic and microscopic tumour) and were followed closely after surgical resection. Of these, C008, B013 and C004 had HCC recurrence at 3.6 months, 6.3 months and 14 months after surgery respectively (**Fig 1B**). All 3 patients with tumor recurrence had unfavourable clinic-pathological features that predisposed to early recurrence, namely tumour size > 5 cm (11.3 cm, 6.7 cm, 6.9 cm respectively), presence of satellite nodules (C008, B013), high TNM stage (IIIA, IIIA, IB) respectively), and alfa-fetoprotein > 400 ng/mL (C004). All 3 tumours were Edmonson Grade 2 and were positive for chronic viral hepatitis B infection. Interestingly, patient A009, who was recurrence-free at 57 months post-surgery also presented with unfavourable clinic-pathological features. This patient was hepatitis B/C virus-negative, had 7.6 cm Edmonson grade 3 HCC with microvascular invasion, and an AFP level of 2203 ng/mL at the time of resection.

### Cell type annotation in multi-sector HCC tumours reveal widespread differences in abundance of cell types that comprise the tumour ecosystem

To determine the abundance of tumour epithelial, stromal, and immune cells in the different tumour sectors, we employed the CellTrek package [28] to annotate the Visium spatial spots using a single cell RNA sequencing data from a recent study by Liu *et al* [29]. This analysis revealed marked heterogeneity in the abundance of cell types that comprise each tumour from a patient, as well as between patients (**Fig 2**). For instance, hepatocytes comprised 35% of the tissue in sector 1 and only 7% in sector 3 of the same tumour (**Fig 2B**). Variabilities in other cell types such as myeloid (17 – 33%), fibroblasts (5 – 12%), endothelial (10.9 – 26.3%) and CD8 T cells (9 – 17%) were also apparent. In addition, we observed that certain cell types, such as regulatory CD4 T cells (0.9 – 3.5%) and exhausted CD8 T cells (0 – 2.7%) comprised a smaller makeup of the entire tumour across patients. These findings suggest that sampling can alter the readout of cellular ecosystem, with implications for the design, execution, and interpretation of single cell studies.

**Figure 2.**
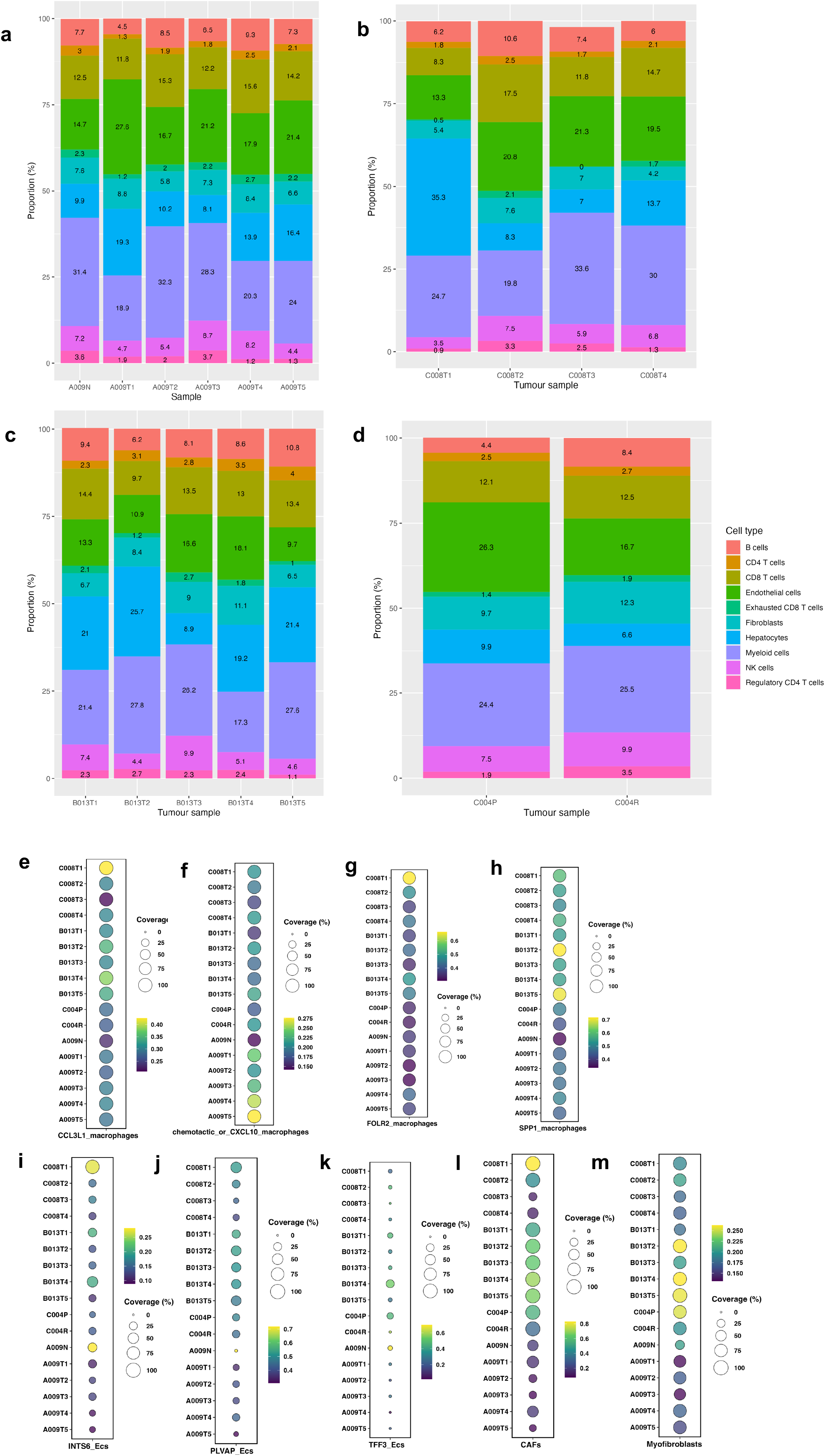
Cellular ecosystem of multi-sector HCC. (a-d) Cell types in each spot of the Visium slide were deconvoluted using the CellTrek package using single cell annotation data from Liu et al. The percentages of each cell type in each sector are depicted as a cumulative bar plot. (e-h) Distribution of macrophage subsets across tumour sectors. (i-k) Distribution of endothelial cell subsets across tumour sectors. (l,m) Distribution of fibroblast subsets across tumour sectors.

Since we observed sizeable variations in the myeloid, fibroblast and endothelial cell populations, we further analysed for subpopulations of these cell types using the annotation provided by Liu *et al*. This analysis revealed that *CXCL10*^+^ macrophage population were enriched in tumour sectors from patient A009 (**Fig 2F**), while cancer associated fibroblasts (CAF) and myofibroblasts were enriched in patients B013, C008 and C004 that experienced disease recurrence (**Fig 2L, M**). It is also worthwhile pointing out that the primary tumour from patient C004 had a higher abundance of both CAFs and myofibroblasts, compared to the secondary tumour. We further annotated the endothelial cells into three populations and observed that there was a trend for lower *INTS6^+^* and *PLVAP*^+^, but not *TFF3*^+^ endothelial cells in patient A009, relative to patients B013, C008 and C004 (**Fig 2I-K**). Altogether, these findings suggest differences in the stromal population of the tumour across geographically distinct sectors associated with disease relapse.

### Gene expression clustering to define intra- and intertumoural heterogeneity

To gain insights into intra- and intertumour molecular heterogeneity present in our patient cohort, we performed gene expression clustering to determine the (i) number of clusters present within each tumour sector and (ii) common and unique clusters within each tumour sector per. We hypothesised that these comparisons may provide a spatial context to gene expression profiles of the tumour ecosystem associated with earlier or later recurrence.

To perform gene expression clustering, we utilised the Squidpy package [30]. As shown in Fig 3, we observed remarkable spatial gene expression heterogeneity across sectors per patient, indicating high levels of intra- and inter-tumour transcriptional diversity across the four patients. For all the sectors analysed, there were between 7-13 gene expression clusters annotated. Interestingly, we also noted that for some sectors where we performed consecutive sections onto the gene expression slide, there were observable changes in gene expression clustering between the two sections that were 14 µm apart, suggesting that consecutive sections may capture transcriptional nuances associated with tissue depth. To explore how the gene expression clusters may be related, we performed hierarchical cluster analysis using the top 5 most variable features per gene expression cluster per tumour sector (**Supp Fig 1**). Interestingly, we observed that the spatial gene expression clusters tend to be located closer to one another on the dendrogram tree, highlighting potential relationship/s among these genes. Overall, the differentially expressed genes in each cluster depicted in the heatmap were associated with specific cell types. For instance, cluster 0 of C008 sector 1 include genes such as *ALB*, *TTR*, and *APOA1*, which represent hepatocytes, cluster 4 (*CD74*, *C1QA*, *C1QB*) represents macrophages, and cluster 6 (*ACTA2*, *TAGLN*, *MCAM*) represents smooth muscle or endothelial cells. Interestingly, cluster 2 (*SAA1*, *SAA2*, *CRP*) represent hepatocytes that express these genes in this patient as well as B013, both of whom experienced recurrence, but not patient A009, who had no disease recurrence. Consistent with this observation, elevated *SAA1* and *SAA2* expression were also observed in intrahepatic cholangiocarcinoma with poor prognosis in a separate study (https://doi.org/10.1101/2021.10.21.465135). A summary of the spatial gene expression and heatmap clusters from other tumour sectors from all patients is provided in Supp Fig 1. Furthermore, we sought pathology evaluation of our H&E images from all the tumour sectors analysed and found very high concordance between our molecular annotation and the broad annotation provided by the pathologist. For instance, in patient B013, there was good agreement in regions identified with tumour cells in most sectors. There were regions identified as fibrotic by the pathologist that was not reflected well in our annotation, which could be due to poor permeabilization hence low RNA yield to ascertain cell types in these regions. In patient C008, tumour cell coverage and fibrotic regions associated with fibroblasts were concordant between our annotation and that of the pathologist. The primary and secondary tumour sectors from patient C004 had relatively low tumour cells, and there was modest concordance in fibroblastic regions.

**Figure 3.**
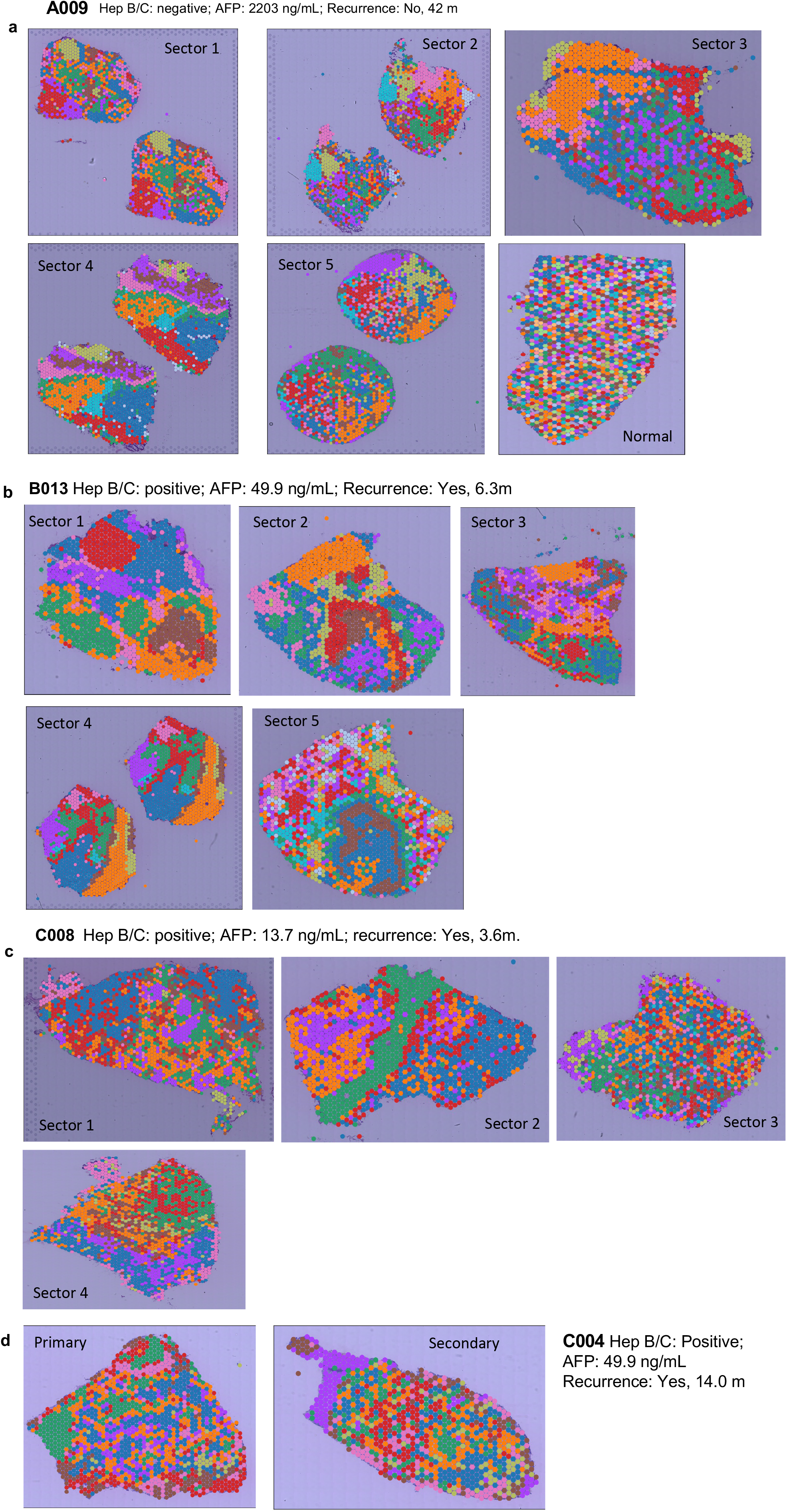
Gene expression clustering reveals widespread transcriptional heterogeneity within and between tumour sectors. (a-d) Gene expression clustering was performed using the Squidpy package. Colors indicate gene expression clusters for each tumour sector and the number of clusters per sector are denoted on the right.

We then asked whether there is significant enrichment in signalling pathways and biological processes between spatial tumour sectors between patients with and without disease recurrence. We first identified differentially expressed genes (DEG) in each cluster and performed pre-ranked gene set enrichment analysis (GSEA) on the DEGs in each gene expression cluster using the HALLMARK database. In patient A009 with no relapse at 46 months of follow up, we observed a trend for increased enrichment of genesets associated with immune and inflammation signalling (e.g., ‘TNFA Signalling via NFKB’, ‘IL2-STAT5 signalling’, ‘Interferon Gamma Response’) across most tumour sectors (**Fig 4A**). However, similar genesets were mostly significantly downregulated in patients B013, C008 and C004, who experienced disease recurrence (**Fig 4B-D**). Altogether, the GSEA suggests that primary tumours with higher levels of immune/inflammatory signalling may be associated with better disease outcome.

**Figure 4.**
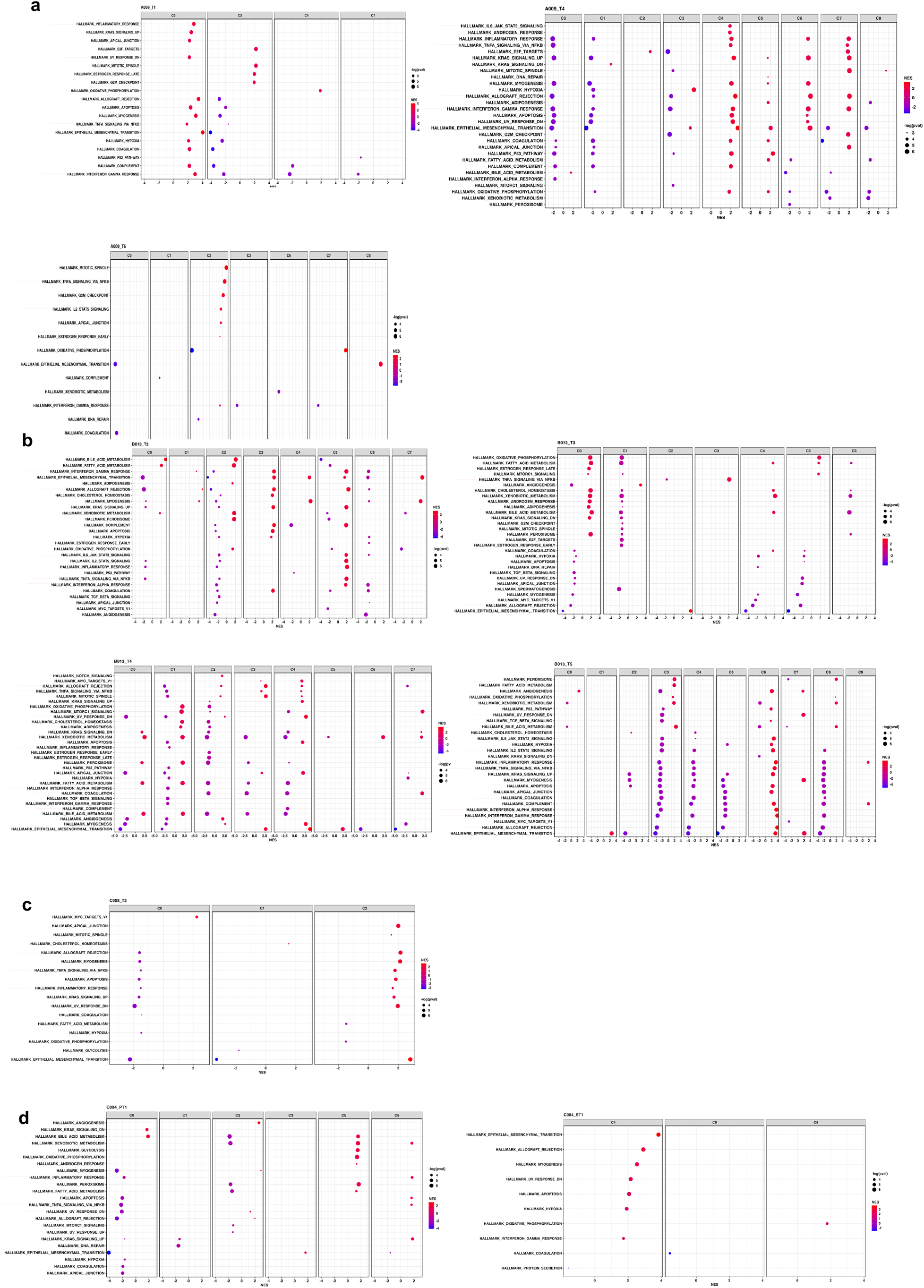
Geneset enrichment analysis of differentially expressed genes in each gene expression cluster. (a) Genesets involved in immune and inflammation signalling are upregulated (indicated in red arrow) in patient A009. (b-d) Most tumour sectors from patients B013, C008 and C004 demonstrate significant reduction in enrichment scores of immune/inflammation signalling.

### Ligand-receptor interaction analysis in primary tumours from patients that do or do not recur

To determine the spatial molecular interactions between the different sectors in recurrent and non-recurrent tumours, we performed ligand-receptor analysis using the Squidpy package [30]. This analysis predicts interactions between the different gene expression clusters in each sector. We noted that apolipoprotein-mediated interactions (e.g., *APOA, APOC*, *APOE*) were highly prominent across many sectors in all patients, consistent with the hepatic identity of these tissues (**Fig 5**).

**Figure 5.**
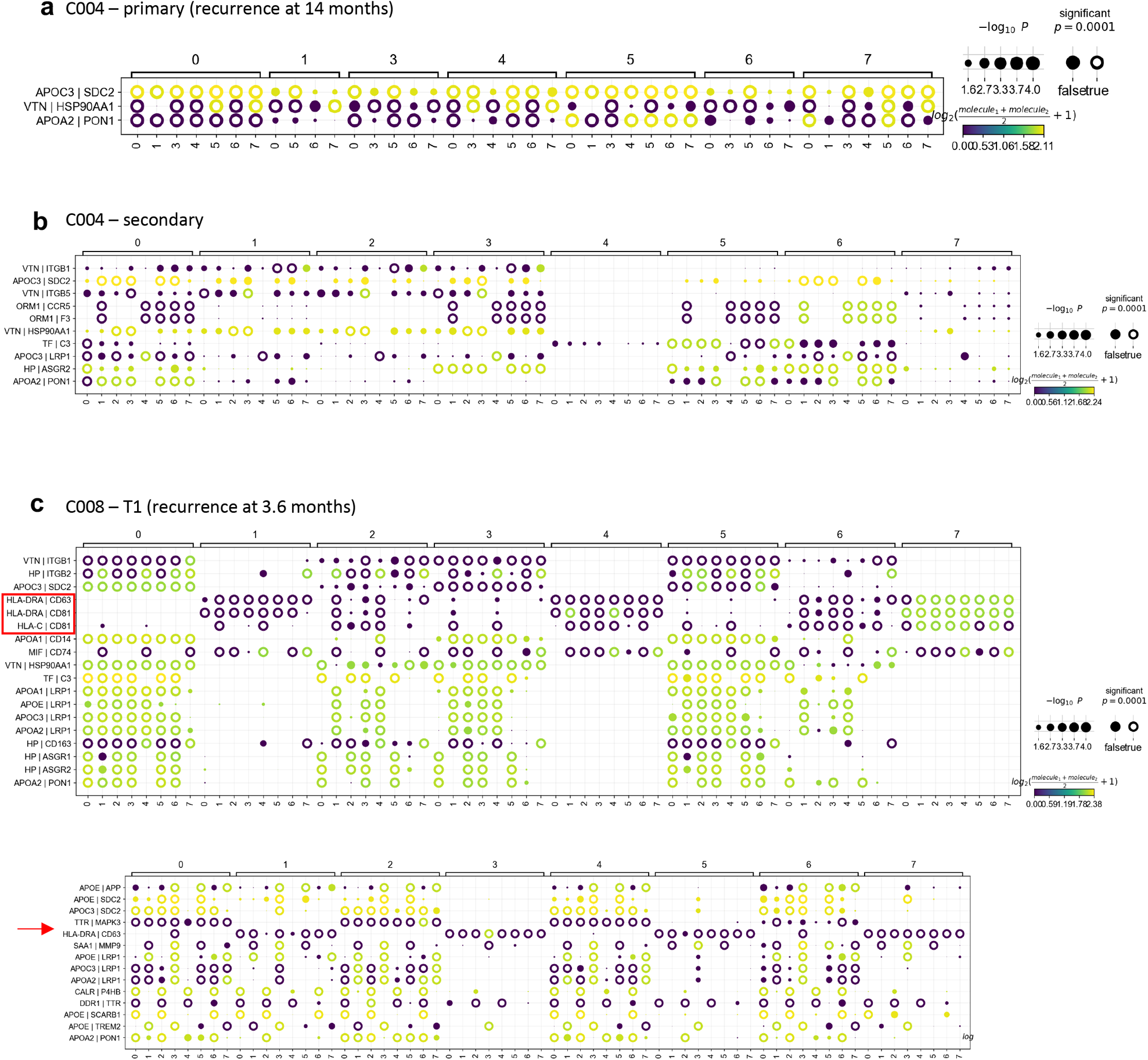
Ligand-receptor interaction analysis identifies enrichment in HLA-mediated interactions in tumour sectors from patients with earlier recurrence. Ligand receptor analysis was performed for between each gene expression cluster for all tumour sectors. Patient C004 primary, C004 secondary, C008 sector 1, and B013 sector 1 are provided as representative examples. Colours and size of circles indicate predicted strength and statistical significance of interaction.

We focused on identifying interactions that were enriched in tumours from patients that eventually recurred, which may shed light on the biology associated with disease relapse. We observed that class I HLA-C and class II HLA-DRA mediated interactions were significantly enriched in tumour sectors associated with earlier recurrence (patients C008 and B013) (**Fig 5C** & **D**), compared to later recurrence at 14 months for patient C004 (**Fig 5A** & **B**). These include HLA-DRA – CD63, HLA-DRA – CD81, HLA-DRA – CTSS and HLA-DRA – CTSL (indicated with red arrows). Furthermore, not all tumour sectors from patients that recurred exhibited the HLA-interactions, suggesting that only a small subpopulation of spatially distinct cellular communication may be involved in recurrence. For instance, in patient B013, HLA-DRA-mediated interactions were only observed in sector 1, in gene expression clusters 0, 1, 2, 4 and 7 and sector 4, in gene expression cluster 0, 5 and 7 (**Fig 5D**). In patient C008, HLA-mediated interactions were observed in gene expression clusters 1, 4, 6 and 7 of sector 1, but not sectors 2 or 3 (**Fig 5C**). Importantly, these interactions were not observed in five independent tumour sectors from patient A009 with no disease recurrence. However, it is worth mentioning that the three patients with recurrence were hepatitis-positive, which may have a role in the enrichment of HLA-mediated interactions observed, relative to patient A009, who was hepatitis-negative.

### Tumour-stroma neighbourhood analysis reveals an enrichment of *TFF3^+^/INT6^+^/PLVAP^+^*endothelial cell proximity to tumour and cancer-associated fibroblasts in patients with recurrence

Next, we sought to understand how the spatial organisation of cell types differ between primary tumours in patients with or without disease relapse. To this end, we employed the CellTrek package [28] and utilised the single cell RNA sequencing data and annotation from a recent study reported by Liu et al. [29] to deconvolute the Visium spatial spots. We first explored the cellular neighbourhoods of hepatocellular carcinoma cells and observed a strong enrichment for *TFF3*^+^/*INT6*^+^/*PLVAP*^+^ endothelial cell in proximity of HCC cells in the primary tumours of C004, C008 and B013, but not patient A009 who remained disease-free after 46 months post-surgery (**Fig 6**). Next, we assessed the cell types that were spatially co-localised in proliferating HCC cells and observed that the primary tumours from the three patients who had disease recurrence were enriched for *SPP1*+ macrophages and proliferating endothelial cells, which was not observed in tumour sectors from patient A009. Altogether, these data demonstrate the differential spatial proximity of specific subtypes of endothelial cells and macrophages with tumour hepatocytes in primary tumours in patients with a history of disease relapse. Cancer-associated fibroblast (CAF) has been proposed to orchestrate crosstalk with tumour epithelial and stromal cells to foster a microenvironment that promotes progression, invasion, and metastasis [31]. Therefore, we applied the neighbourhood analysis to investigate the cellular proximities associated with CAFs. Interestingly, we also observed *INTS6^+^* endothelial cell as a common neighbour for CAFs in the C004, C008 and B013 tumour sectors (**Fig 6B-D**), while *PLVAP*^+^ endothelial cell was predicted to be in proximity with CAFs in patient A009 (**Fig 6A**).

**Figure 6.**
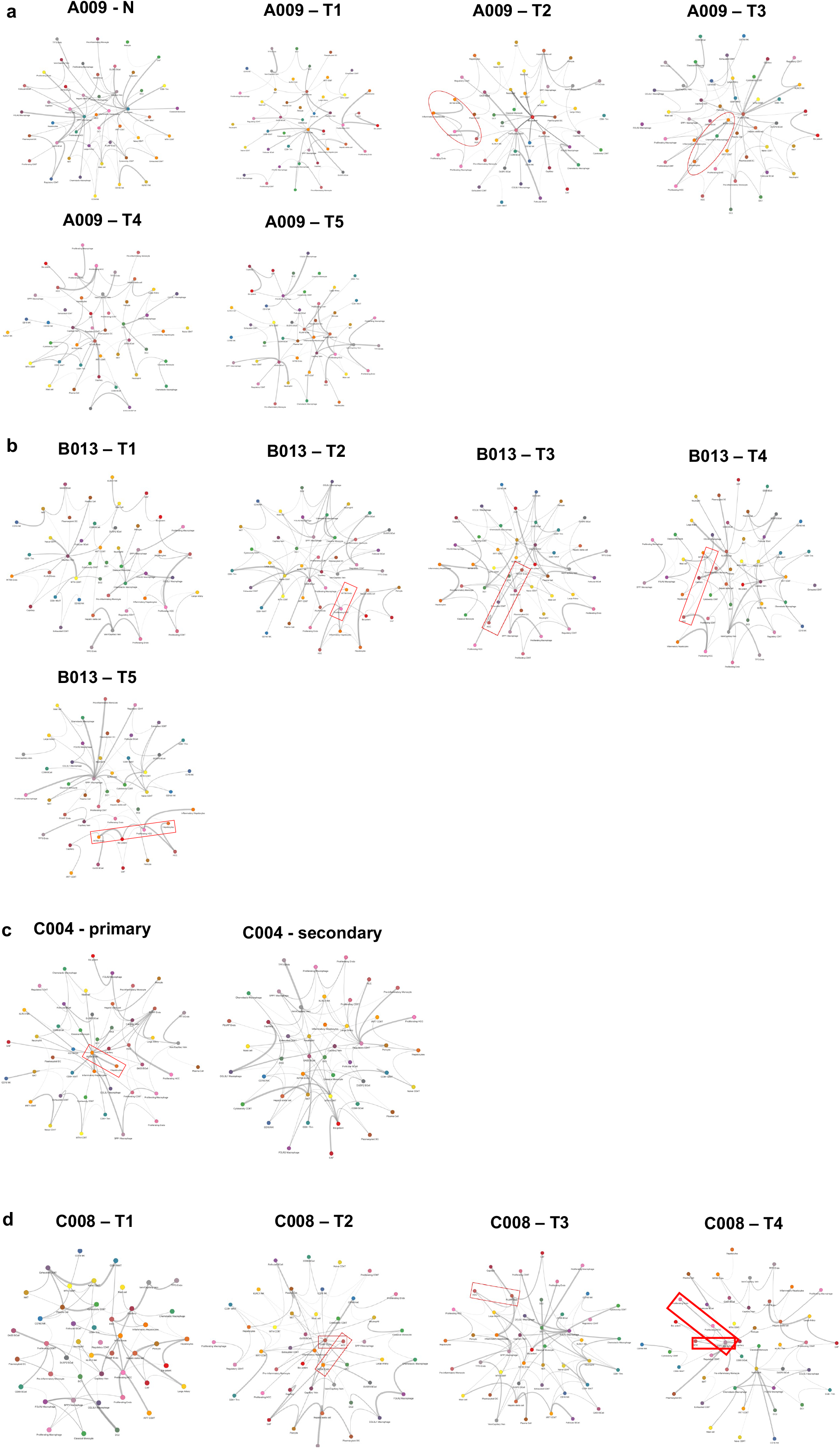
Cellular neighbourhood analysis identifies an enrichment for *INTS6*+ endothelial cells and *SPP1*+ macrophage in close proximity to hepatocytes in patients with recurrence. Selected tumour sectors where hepatocytes are in proximity with endothelial cells are highlight in red.

Since we observed an enrichment for *INTS6^+^* endothelial cell and *SPP1^+^* macrophages in primary tumours of patients who experienced recurrence, we further probed whether *INTS6* or *SPP1* expression was associated with progression-free (PFS) and overall (OS) survival using bulk RNA seq and follow up data from a local cohort of 91 patients previously reported (Supplementary Table 1 on their demographics). High expression of *SPP1* was significantly correlated with poorer prognosis in our cohort (**Supp Fig 2**), and although a similar trend was observed for *INTS6*, the survival curves did not reach statistical significance (**Fig 7A** & **B**). Since *SPP1* association with prognosis has already been reported [32], we turned our focus to *INTS6^+^* endothelial cells and sought to understand the gene expression differences in the *INTS6*+ endothelial, relative to the other endothelial cell population. To this end, we utilised the single cell RNA seq dataset from Liu *et al* to perform differential expression (DEG) analysis (**Fig 7C - E**). There were 516 genes that were upregulated, and 256 genes downregulated in the *INTS6*^+^ positive compared to all other endothelial cells (Supplementary Table 2). Importantly, we noted that a selected number of these genes were strongly upregulated only in *INTS6*^+^ endothelial cells, but not other immune and stromal cell types analysed, suggesting a gene signature associated with this specific cell type (**Fig 7E**). Interestingly, the *PLVAP* gene, which is a key marker of endothelial cells [33], was downregulated approximately two-fold in *INST6*^+^ endothelial cell, suggesting that the latter may represent a unique population of endothelial cell. To determine the pathways that were over-represented among the differentially expressed genes, we performed pathway enrichement analysis using the MSigDB database (**Fig 7F**). We focused on genes with log2 fold change of at least ± 0.5 and observed that pathways enriched in the overexpressed genes included ‘TNF Signaling’, ‘Hypoxia’, ‘Complement’, ‘Androgen Response’, and ‘Apoptosis’, while pathways that were enriched in genes downregulated included ‘Hypoxia’ and ‘Epithelial-to-Mesenchymal Transition’. Thus, these findings suggest a potential inflammatory-hypoxia interaction enriched in the *INST6*+ endothelial cells. Next, since we observed the *INTS6^+^* endothelial cell in close proximity to tumour cells, we performed ligand-receptor interaction analysis to determine potential mediators of cellular communication associated with the tumours that relapse. We identified several significant interactions, including *ANGPLT4 – SDC1, COL4A1/2-SDC1,* and *SPP1-ITGA5/ITGB1*. The SDC1 gene has been implicated in various oncogenic processes, thereby supporting its enrichment in the interactions identified [34, 35].

**Figure 7.**
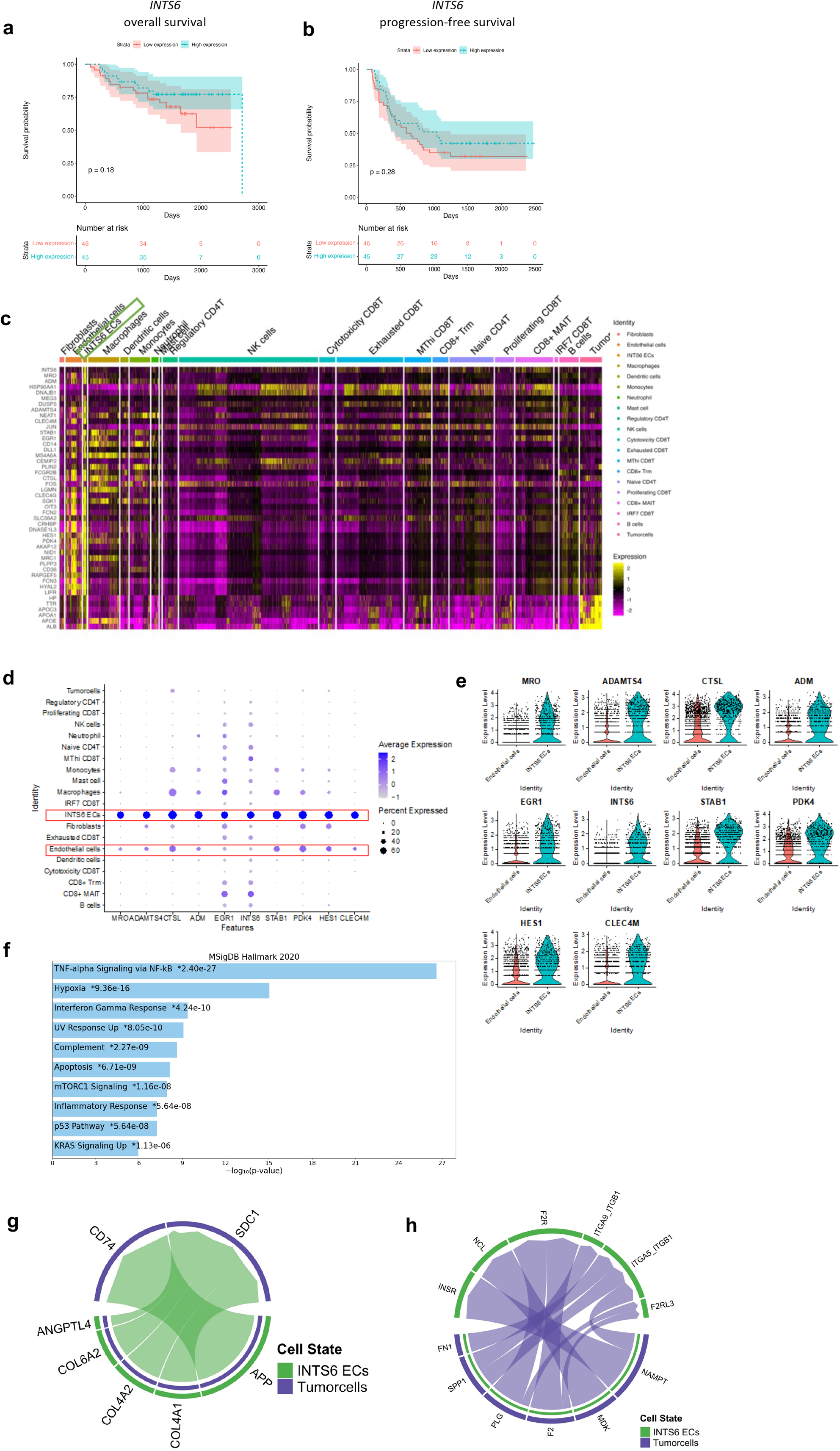
*INTS6+* endothelial cell gene signature. (a,b) High *INTS6* expression reveal a non-significant trend association with overall and progression-free survival. (c) Top 50 genes differentially expressed in between *INTS6+* endothelial cell compared to other cell types. (d,e) Dot plots and violin plots to demonstrate that the unique gene expression signature associated with *INST6^+^* endothelial cells. (f) Pathways enriched in genes upregulated in *INST6^+^* endothelial cells, relative to other endothelial cell population.

### Spatial heterogeneity of immuno-related signatures and targets across tumour sectors

The IMbrave150 study demonstrated improved survival for patients that received atezolizumab (anti-PD-L1) and bevacizumab (anti-VEGFA) compared to sorafenib [36]. Based on the findings from this trial, combination treatment with both agents has been approved as the first-line systemic treatment for unresectable disease. However, treatment resistance remains a clinical challenge and the molecular mechanisms is poorly understood. We explored the spatial expression of two published immune-related signatures in our patient cohort [20, 37]. The 50-gene tertiary lymphoid structure signature (TLS50) marks regions of the tumour with immune cell aggregates and correlated with favourable response to immunotherapy [38]. We mapped the TLS50 signature on our tumour sectors and observed that the signature expression was heterogeneous and formed pockets of regions when expressed, consistent with the TLS being aggregates of immune cells clustered together (**Fig 8A-D**). In this cohort, patient B013 had the highest expression of TLS50, compared to other patients (**Fig 8C**). Interestingly, patient A009 with no recurrence at 46 months of follow up post-surgery, exhibited very low TLS50 signature expression across all sectors (**Fig 8A**) while in patient C004, we noted that the TLS50 signature was low in both the primary and secondary tumour sectors (**Fig 8D**).

**Figure 8.**
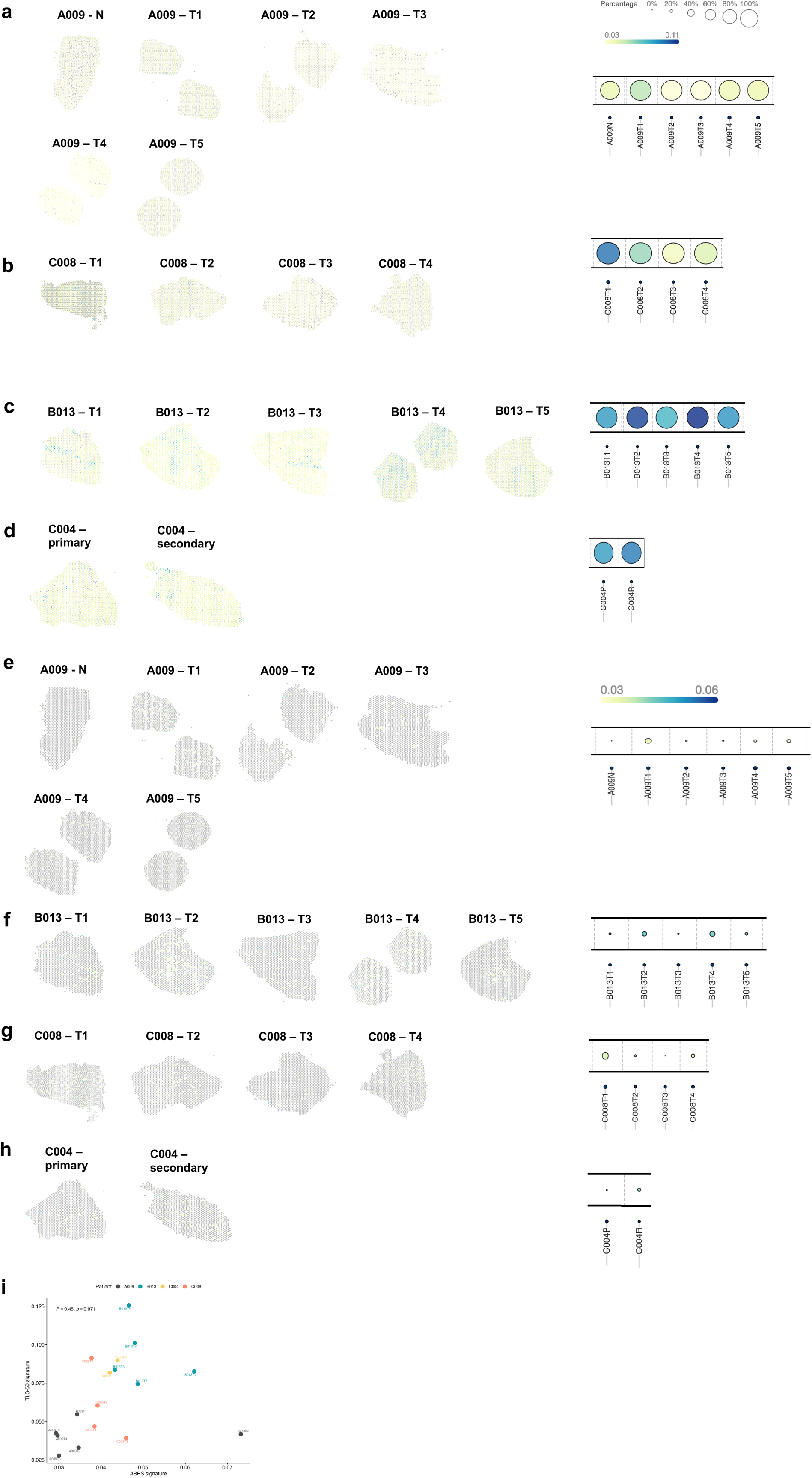
Spatial heterogeneity of tertiary lymphoid structures and the ABR signatures across tumour sectors. (a-d) Spatial expression profile of the TLS50 signature in patients A009, B013, C008, C004. (e-h) Spatial expression profile of the ABR signature in patients A009, B013, C008, C004. Summary of each tumour sector is indicated as a bubbleplot on the right and the respective scales indicated. (i) Correlation analysis between the TLS50 and ABR signatures.

A recent study utilised patient samples available from the IMbrave150 [36] and GO30140 [39] trials to investigate the molecular features associated with response to atezolizumab, bevacizumab, or sorafenib and reported a 10-gene atezolizumab + bevacizumab response (ABR) signature [37]. Patient tumours with high expression of the ABR signature demonstrated better progression-free and overall survival to atezolizumab + bevacizumab treatment, compared to high ABR expression in tumours treated with sorafenib only. However, this signature was derived from the bulk tumour, which precludes understanding the spatial distribution of these 10 genes. Thus, we analysed the spatial expression of the ABR signature in our patient cohort. In general, we observed low signature expression across the tumour sectors, although there was only some degree of variability observed between sectors (**Fig 8E-H**). Since both signatures are associated with response to immune checkpoint blockade, we performed a correlation analysis and noted a trend for a positive relationship between the TLR50 and ABR signatures (**Fig 8I**).

One treatment modality being explored in the treatment of HCC is the use of chimeric antigen receptor T cells (CAR-T) to target antigens expressed on tumour cells [40]. While clinical studies are ongoing to evaluate CAR-T response, several mechanisms of resistance have been proposed, including modulation of target expression [41]. Thus, we explored the spatial expression profile of two HCC tumour antigens, AFP and GPC3, where CAR-T cells have been developed and patients are undergoing recruitment for clinical investigation (NCT04121273, NCT02905188, NCT03198546, NCT03884751) (**Supp Fig 3**). Patient A009, which had a serum AFP of 2203 ng/mL, showed consistently high expression of *AFP* in all sectors analysed, with negligible expression in normal liver tissue. In contrast, patient C008, which had a serum AFP of 13.7 ng/mL, exhibited low *AFP* expression across all sectors, while patient B013, showed heterogeneous expression of the *AFP* transcript across all sectors (serum AFP 49.9 ng/mL). Interestingly, while patient C004 had high serum AFP levels (999 ng/mL), the one primary tumour sector demonstrated low levels of the *AFP* mRNA. Thus, these findings suggest that expression of two key HCC tumour antigens may vary and argues that treatment decision should be guided by comprehensive profiling of targeted antigens at spatial resolution, rather than bulk tumour tissues.

## Discussion

In this study, we sought to provide a spatial cellular atlas of hepatocellular carcinoma in patients with or without disease relapse after surgery. Since HCC has a bimodal distribution of recurrence over time [4], we focused on patients that exhibited recurrence in less than 2 years, which has been proposed to likely be related to the primary tumour micrometastasis.

In the cell type distribution analysis, we observed variabilities in most cell types annotated. In particular, the myeloid, fibroblast, and endothelial cell populations were variable across tumour sectors. Thus, this argues for study designs that consider profiling multiple regions of the tumour as single-site sampling may not fully capture the cellular complexity, and thus, biology of the tumour [42, 43]. We further explored the three cell populations mentioned and observed that the *CXCL10*^+^ macrophages were enriched in the tumour sectors of patient A009 who did not experience recurrence while the *INTS6*^+^ and *PLVAP*^+^ endothelial cells were more abundant in patients B013, C008 and C004. Several studies have demonstrated that *CXCL10*^+^ macrophages are necessary for inducing anti-tumour inflammation and response to chemotherapy and immune checkpoint blockade [44–46]. In contrast, how *INTS6*^+^ endothelial cells function in the context of HCC recurrence is poorly understood. However, the *PLVAP*^+^ endothelial cells identified in our analysis has been proposed as a therapeutic target and was recently described as part of an oncofetal niche that forms an immunosuppressive environment in HCC [17, 47], thereby suggesting that the *INTS6*^+^ population identified is a novel association. The *INST6* gene encodes a putative RNA helicase and its role in endothelial cell function is poorly described, thus, mechanistic experiments are required to better understand how this endothelial cell subtype features in HCC biology. Our cellular neighbourhood analysis revealed recurrent proximities of *INTS6*^+^ endothelial cell and *SPP1*^+^ macrophage with tumour hepatocytes and CAFs. Importantly, recent studies have demonstrated for a role of the *SPP1*^+^ macrophage as part of a stromal signature that predicts poor recurrence and associated with poor response to lenvatinib/sorafenib [48, 49]. Interestingly, our ligand-receptor interaction analysis identified *SPP1* interacted with *CD44* in only one tumour sector from patient B013 who had a recurrence at 6.3 months post-surgery. The *SPP1*-*CD44* interaction was previously reported to be associated with poor prognosis and *SPP1* could induce the polarisation of macrophage towards the M2 state, thereby adopting an anti-inflammatory profile [50]. Based on our data, functional studies that assess the interruption of *SPP1*^+^ macrophage and *INTS6*^+^ endothelial cell may yield novel insights into how spatial interactions of the tumour stroma affects tumour biology.

Our geneset enrichment analysis suggests a trend for increased inflammation and immune-related signalling in patient A009 who did not experience relapse and was lower in the three patients who relapsed. However, we observed heterogeneity in expression of these inflammatory/immune genesets across tumour sectors of each patient, albeit the overall trend stands when assessed across all the sectors. One possible reason for elevated inflammation in the tumour of patient A009 is that the individual developed dermatomyositis as part of a relatively rare paraneoplastic syndrome from HCC, which mounts a systemic immune response. Nevertheless, the excellent prognosis of this patient associated with an inflammatory tumour profile is consistent with the findings of Chew *et al* [51]. Patient B013 was a HepB charrier in the immune-tolerance phase, suggesting a blunted immune response to the virus, consistent with the relatively lower expression of immune/inflammatory genesets in the tumour. Similarly, for patient C004 who was HepB-positive and received pre-surgery anti-viral treatment, the viral burden was probably not completely cleared. Nevertheless, findings from our gene set enrichment analysis and the clinical information of the HepB-positive patients agree with a recent study that suggests that HepB infected patients demonstrate a tumour immunosuppressive state [52]. On a technical note, given that this enrichment analysis is highly informative of tumour biology, these findings further reiterate the importance of multi-region sampling as the insufficient assessment of tumour regions may under- or overestimate specific pathways of interest.

Our exploration of published gene expression signatures associated with tumour immunology and response to immune checkpoint blockade agent revealed marked heterogeneity across tumour sectors. This suggests that while regions with high expression of the TLS50 and/or ABR signatures may respond well to immune checkpoint blockade, such treatment will also induce the positive selection of population of cells with low signature expression, leading to treatment resistance. We noted that patient B013 had relatively high ABR signature expression across all tumour sectors, compared to patient A009, who had low signature expression, which raises the question worth addressing in future clinical studies: instead of how high, how low an expression of the ABR/TLS50 signatures is required to achieve durable response? Lastly, we also observed a modest positive correlation between the ABR and TLS50 signatures, consistent with the role for tertiary lymphoid structures in promoting better response to immune checkpoint blockade. This correlation also raises the possibility of constructing a model that incorporates the most significantly predictive features of each model to establish a better predictor of treatment response. In conclusion, our study has defined the multi-region spatial transcriptomic landscape of HCC tumours with and without recurrence post-surgery. This dataset will provide a valuable resource for the community to further explore pathways on interest, and the associated cellular neighbourhoods and interactions to better understand HCC biology.

