## Supplementary Table 1 for "Multi-region spatial transcriptome analysis reveals cellular networks and pathways associated with hepatocellular carcinoma recurrence"

| Patient ID | Gender | Race | Child's Pugh | Age (at surgery) | TNM stage | Edmonson grade | Microvascular invasion | HBV Status | HCV Status | AFP (ng/ml) |
|---|---|---|---|---|---|---|---|---|---|---|
| HEP0026 | Male | Indonesian Chinese | A | 72 | II | 3 | yes | 0 | 0 | 348 |
| HEP0098 | Male | Indonesian | A | 57 | IB | 2 | no | B | 0 | 28 |
| HEP0174 | Male | Chinese | A | 66 | IB | 2 | no | B | 0 | 7.2 |
| HEP0034 | Male | Chinese | A | 75 | IB | 3 | no | B | 0 | 3.9 |
| HEP0076 | Female | Malay | A | 68 | II | 3 | yes | 0 | 0 | 422 |
| HEP0152 | Male | Chinese | A | 60 | IIIA | 3 | yes | 0 | 0 | 17349 |
| HEP0186 | Male | Chinese | A | 63 | IB | 2 | no | B | 0 | NA |
| HEP0193 | Male | Chinese | A | 63 | IB | 2 | no | 0 | 0 | 8.6 |
| HEP0200 | Male | Chinese | A | 87 | IB | 3 | no | 0 | 0 | 10.4 |
| HEP0206 | Female | Chinese | A | 50 | IB | 3 | no | B | 0 | NA |
| HEP0207 | Male | Chinese | A | 68 | IB | 2 | no | B | 0 | 3.1 |
| HEP0209 | Female | Chinese | A | 54 | IB | 2 | no | B | 0 | 4.6 |
| HEP0229 | Male | Chinese | A | 80 | IIIB | 3 | yes | 0 | 0 | 5.8 |
| HEP0235 | Male | Chinese | A | 53 | IB | 2 | no | B | 0 | 7.1 |
| HEP0241 | Male | Indian | A | 61 | IB | 2 | no | 0 | 0 | NA |
| HEP0262 | Male | Chinese | A | 63 | IIIA | 3 | yes | B | 0 | 31.5 |
| HEP0269 | Male | Sikh | A | 65 | IB | 1 | no | 0 | 0 | 1.8 |
| HEP0270 | Male | Chinese | A | 68 | IA | 3 | no | B | 0 | 3 |
| A001 | Male | Chinese | A | 66 | IA | 4 | no | 0 | 0 | 6.7 |
| A004 | Male | Chinese | A | 77 | II | 3 | yes | B | 0 | 38 |
| A005 | Male | Chinese | A | 65 | IA | 3 | no | B | 0 | 69.3 |
| HEP0247 | Female | Chinese | A | 82 | II | 3 | yes | 0 | 0 | 27.3 |
| HEP0264 | Male | Malay | A | 64 | II | 2 | yes | 0 | 0 | 20.1 |
| HEP0276 | Male | Chinese | A | 76 | IB | 3 | no | 0 | 0 | 3.2 |
| HEP0277 | Male | Chinese | A | 70 | II | 3 | yes | B | 0 | NA |
| HEP0319 | Male | Indonesian | A | 69 | IIIB | 3 | yes | 0 | 0 | 14483 |
| HEP0321 | Male | Chinese | A | 70 | IB | 2 | no | 0 | 0 | 2.1 |
| HEP0356 | Male | Others | A | 76 | II | 2 | yes | 0 | C | 106 |
| HEP0261 | Male | Malay | A | 57 | IB | 3 | no | 0 | C | NA |

| | | | | | | | | | | |
|---|---|---|---|---|---|---|---|---|---|---|
| A009 | Female | Chinese | A | 47 | II | 3 | yes | 0 | 0 | 2203 |
| HEP0507 | Male | Others | A | 55 | II | 3 | yes | B | 0 | 1.9 |
| HEP0549 | Male | Chinese | A | 70 | IB | 2 | no | 0 | 0 | 3.14 |
| A011 | Male | Chinese | A | 71 | IB | 2 | no | B | 0 | 4.9 |
| B002 | Male | Chinese | A | 52 | IB | 2 | No | B | 0 | 4.7 |
| B003 | Male | Chinese | A | 60 | IIIB | 3 | yes | 0 | 0 | 1.9 |
| B004 | Male | Chinese | A | 52 | IB | 3 | No | B | 0 | 13.5 |
| B006 | Female | Chinese | A | 67 | IB | 3 | No | B | 0 | 8592 |
| B007 | Male | Chinese | A | 68 | IA | 2 | No | 0 | 0 | 56.2 |
| B008 | Male | Chinese | A | 78 | IIIB | 3 | yes | B | 0 | 36.6 |
| B009 | Male | Malay | A | 67 | IIIB | 3 | yes | B | 0 | >60500 |
| B010 | Female | Chinese | A | 74 | IB | 1 | No | 0 | 0 | 5.4 |
| B011 | Male | Chinese | A | 60 | II | 3 | yes | B | 0 | 3.5 |
| B012 | Male | Chinese | A | 62 | IB | 1 | No | B | 0 | 4.2 |
| B013 | Male | Chinese | A | 57 | IIIA | 2 | No | B | 0 | 49.9 |
| B014 | Male | Chinese | A | 73 | II | 2 | yes | B | 0 | 1.5 |
| B015 | Male | Chinese | A | 63 | IB | 2 | No | B | 0 | 6.1 |
| B017 | Male | Chinese | A | 75 | II | 3 | yes | B | 0 | 504 |
| B018 | Male | Chinese | A | 46 | IB | 3 | No | B | 0 | 657 |
| B019 | Male | Chinese | A | 63 | IIIA | 2 | yes | B | 0 | 62 |
| B020 | Male | Chinese | A | 80 | II | 1 | yes | 0 | 0 | 2.5 |
| B021 | Male | Chinese | A | 67 | IIIA | 2 | yes | B | 0 | 9.6 |
| B022 | Female | Chinese | A | 71 | II | 3 | yes | B | 0 | 375 |
| B023 | Male | Malay | A | 64 | II | 3 | No | 0 | 0 | 6.3 |
| B024 | Male | Chinese | A | 76 | IB | 2 | No | B | 0 | 1.9 |
| B025 | Male | Chinese | A | 73 | IB | 2 | No | 0 | 0 | 1029 |
| B026 | Female | Chinese | A | 70 | II | 3 | yes | 0 | 0 | 3121 |
| C002 | Male | Chinese | A | 69 | IIIA | 2 | No | 0 | 0 | 1069 |
| C003 | Male | Chinese | A | 72 | II | 2 | No | B | 0 | 3 |
| C004 | Female | Chinese | A | 69 | IB | 2 | No | B | 0 | 999.99 |
| C006 | Male | Chinese | A | 70 | IB | 2 | No | B | 0 | 2.5 |
| C008 | Male | Chinese | A | 75 | IIIA | 2 | No | B | 0 | 13.7 |

| | | | | | | | | | | |
|---|---|---|---|---|---|---|---|---|---|---|
| C011 | Male | Indian | A | 70 | II | 2 | No | B | 0 | 6.5 |
| C012 | Male | Chinese | A | 71 | II | 3 | yes | 0 | 0 | 54458 |
| C014 | Male | Chinese | A | 60 | II | 2 | yes | B | 0 | 2.7 |
| C017 | Male | Chinese | A | 65 | IB | 2 | No | B | 0 | NA |
| D002 | Male | Chinese | A | 70 | IB | 1 | No | B | 0 | 1 |
| D006 | Male | Chinese | A | 62 | II | 2 | No | 0 | C | 134.939759 |
| D007 | Male | Indian | A | 22 | IB | 2 | No | B | 0 | 1245.78313 |
| D008 | Male | Chinese | A | 66 | IB | 1 | No | 0 | 0 | 9.63855422 |
| D010 | Male | Chinese | A | 54 | II | 2 | yes | B | 0 | 16.8674699 |
| D011 | Female | Chinese | A | 73 | II | 3 | yes | B | 0 | 102.409639 |
| D012 | Male | Chinese | A | 74 | II | 4 | yes | B | 0 | 425.301205 |
| E001 | Female | Thai | A | 79 | IB | 2 | No | B | 0 | 11304 |
| E002 | Female | Thai | A | 53 | IB | 2 | No | B | 0 | 461.2 |
| E003 | Male | Thai | A | 59 | II | 2 | yes | B | 0 | 1.59 |
| E004 | Male | Thai | A | 67 | II | 2 | No | B | 0 | 831.7 |
| E006 | Male | Thai | A | 57 | IB | 2 | No | B | 0 | 64.53 |
| E008 | Male | Thai | A | 60 | IB | 1 | No | B | 0 | 3.15 |
| E009 | Male | Thai | A | 61 | IB | 1 | No | B | C | 12.17 |
| E010 | Female | Thai | A | 75 | IA | 1 | No | B | 0 | 86.81 |
| E012 | Male | Thai | A | 51 | IB | 2 | No | B | 0 | 925 |
| E014 | Male | Thai | A | 61 | II | 2 | yes | 0 | C | 999.99 |
| E015 | Male | Thai | A | 36 | IB | 1 | No | B | 0 | 4.64 |
| E016 | Female | Thai | A | 57 | IB | 3 | No | B | C | 3512 |
| E017 | Male | Thai | A | 57 | IB | 2 | No | B | 0 | 147 |
| F002 | Male | Filipino | A | 77 | II | 2 | yes | B | 0 | 26.6 |
| F004 | Male | Filipino | A | 53 | II | 2 | yes | B | 0 | NA |
| F005 | Male | Filipino | A | 66 | II | 1 | No | B | 0 | 1.5 |
| F006 | Male | Filipino | A | 77 | II | 3 | yes | B | 0 | 446.4 |
| F008 | Male | Filipino | A | 51 | II | 4 | yes | B | 0 | NA |
| F009 | Male | Chinese | A | 73 | II | 2 | yes | 0 | 0 | 1936.46 |
