## Supplementary Table 2 for "Multi-region spatial transcriptome analysis reveals cellular networks and pathways associated with hepatocellular carcinoma recurrence"

| Gene symbol | p_val | avg_log2FC | pct.1 | pct.2 | p_val_adj |
|---|---|---|---|---|---|
| MGP | 4.76824E-50 | -2.282557381 | 0.13 | 0.423 | 1.1809E-45 |
| VIM | 2.24021E-97 | -1.933377968 | 0.281 | 0.715 | 5.5479E-93 |
| VWF | 7.89428E-37 | -1.588358628 | 0.124 | 0.367 | 1.955E-32 |
| SPARCL1 | 2.97606E-40 | -1.521184258 | 0.051 | 0.301 | 7.3702E-36 |
| S100A6 | 1.61118E-67 | -1.288870661 | 0.265 | 0.621 | 3.9901E-63 |
| PLVAP | 1.59313E-43 | -1.122834259 | 0.073 | 0.346 | 3.9454E-39 |
| IGFBP7 | 7.53588E-31 | -0.983050694 | 0.808 | 0.873 | 1.8663E-26 |
| TMSB10 | 6.6167E-104 | -0.975243899 | 0.992 | 0.998 | 1.639E-99 |
| EMP1 | 5.63984E-21 | -0.972770305 | 0.203 | 0.373 | 1.3967E-16 |
| CLU | 6.64382E-12 | -0.95337661 | 0.109 | 0.222 | 1.6453E-07 |
| SLC9A3R2 | 3.15893E-27 | -0.918125023 | 0.572 | 0.676 | 7.8231E-23 |
| IGFBP3 | 7.00193E-08 | -0.917170662 | 0.116 | 0.198 | 0.00173403 |
| AQP1 | 4.86142E-44 | -0.915669209 | 0.033 | 0.295 | 1.2039E-39 |
| LGALS1 | 2.58876E-28 | -0.882320804 | 0.116 | 0.338 | 6.4111E-24 |
| FN1 | 1.33957E-32 | -0.871831046 | 0.039 | 0.253 | 3.3174E-28 |
| CAV1 | 5.45405E-28 | -0.853650278 | 0.29 | 0.48 | 1.3507E-23 |
| DEPP1 | 7.27954E-13 | -0.851968416 | 0.104 | 0.223 | 1.8028E-08 |
| SRP14 | 1.70705E-50 | -0.84758435 | 0.791 | 0.888 | 4.2275E-46 |
| PECAM1 | 8.35789E-34 | -0.844944654 | 0.488 | 0.624 | 2.0698E-29 |
| PODXL | 1.15632E-35 | -0.81353582 | 0.169 | 0.407 | 2.8636E-31 |
| SPRY1 | 2.91821E-25 | -0.803860505 | 0.183 | 0.379 | 7.227E-21 |
| CRIP2 | 9.16468E-37 | -0.757789036 | 0.315 | 0.528 | 2.2696E-32 |
| RPS28 | 2.32193E-61 | -0.733783317 | 0.975 | 0.998 | 5.7503E-57 |
| CD34 | 1.49995E-47 | -0.731460858 | 0.064 | 0.353 | 3.7146E-43 |
| GSN | 6.89004E-25 | -0.7305419 | 0.299 | 0.477 | 1.7063E-20 |
| RPS18 | 3.62306E-63 | -0.720334642 | 0.915 | 0.993 | 8.9725E-59 |
| S100A10 | 1.92586E-37 | -0.717570516 | 0.719 | 0.807 | 4.7694E-33 |
| ADIRF | 7.37086E-18 | -0.71402642 | 0.118 | 0.267 | 1.8254E-13 |
| PALMD | 9.6443E-49 | -0.712657102 | 0.054 | 0.345 | 2.3884E-44 |
| TMSB4X | 1.36136E-55 | -0.706320602 | 0.998 | 1 | 3.3714E-51 |
| UTRN | 5.16769E-23 | -0.696709076 | 0.363 | 0.516 | 1.2798E-18 |
| LMNA | 1.15943E-20 | -0.690170749 | 0.344 | 0.498 | 2.8713E-16 |
| MTRNR2L12 | 1.1996E-16 | -0.676869555 | 0.89 | 0.813 | 2.9708E-12 |
| RPL8 | 1.34395E-60 | -0.676432805 | 0.816 | 0.966 | 3.3283E-56 |

| | | | | | |
|---|---|---|---|---|---|
| JAG1 | 6.32674E-18 | -0.675278877 | 0.104 | 0.253 | 1.5668E-13 |
| PTMA | 3.21693E-62 | -0.674096485 | 0.941 | 0.994 | 7.9667E-58 |
| RBP7 | 1.08021E-13 | -0.659128804 | 0.102 | 0.225 | 2.6751E-09 |
| CLEC14A | 2.06903E-20 | -0.657565634 | 0.268 | 0.425 | 5.1239E-16 |
| RGCC | 2.95484E-11 | -0.657445669 | 0.102 | 0.216 | 7.3177E-07 |
| RPL36 | 3.12358E-50 | -0.656337705 | 0.941 | 0.984 | 7.7355E-46 |
| RPL34 | 5.57279E-57 | -0.65102548 | 0.93 | 0.99 | 1.3801E-52 |
| RPS6 | 9.71698E-52 | -0.623123092 | 0.876 | 0.982 | 2.4064E-47 |
| ANXA2 | 1.05836E-30 | -0.618893884 | 0.498 | 0.666 | 2.621E-26 |
| RPL32 | 8.63415E-50 | -0.615472492 | 0.926 | 0.99 | 2.1382E-45 |
| STC1 | 6.02723E-18 | -0.615428019 | 0.036 | 0.166 | 1.4926E-13 |
| RPS23 | 1.19237E-52 | -0.607823888 | 0.91 | 0.989 | 2.9529E-48 |
| RPS15 | 6.90866E-56 | -0.605348895 | 0.902 | 0.989 | 1.7109E-51 |
| S100A11 | 3.53818E-32 | -0.599206296 | 0.597 | 0.734 | 8.7623E-28 |
| RPS8 | 4.06285E-51 | -0.598607382 | 0.902 | 0.987 | 1.0062E-46 |
| RPS24 | 2.02083E-54 | -0.598485102 | 0.907 | 0.985 | 5.0046E-50 |
| FKBP1A | 1.35783E-23 | -0.596596543 | 0.664 | 0.729 | 3.3627E-19 |
| FBLN2 | 2.12289E-23 | -0.59641503 | 0.023 | 0.177 | 5.2573E-19 |
| RPL11 | 6.17281E-55 | -0.593675803 | 0.916 | 0.987 | 1.5287E-50 |
| RAMP2 | 2.24863E-22 | -0.593079716 | 0.589 | 0.694 | 5.5687E-18 |
| TAGLN2 | 2.57606E-33 | -0.591092916 | 0.33 | 0.554 | 6.3796E-29 |
| CD9 | 8.1875E-16 | -0.578603489 | 0.322 | 0.473 | 2.0276E-11 |
| RPL3 | 1.12195E-56 | -0.570601194 | 0.854 | 0.981 | 2.7785E-52 |
| RPS3A | 1.16624E-52 | -0.569642362 | 0.811 | 0.968 | 2.8882E-48 |
| RPL26 | 6.31485E-46 | -0.562947864 | 0.809 | 0.966 | 1.5639E-41 |
| RPL29 | 2.80598E-44 | -0.561300623 | 0.791 | 0.955 | 6.949E-40 |
| RPL37 | 3.2926E-49 | -0.559863535 | 0.867 | 0.974 | 8.1541E-45 |
| SERF2 | 1.06315E-39 | -0.55662046 | 0.964 | 0.979 | 2.6329E-35 |
| RPL7A | 1.06329E-44 | -0.552293749 | 0.837 | 0.964 | 2.6332E-40 |
| RPL30 | 9.63953E-44 | -0.543480904 | 0.879 | 0.979 | 2.3872E-39 |
| RPS19 | 1.11815E-44 | -0.54105057 | 0.916 | 0.99 | 2.7691E-40 |
| NACA | 5.0768E-41 | -0.53739478 | 0.741 | 0.913 | 1.2573E-36 |
| PTMS | 5.99652E-25 | -0.532868017 | 0.499 | 0.636 | 1.485E-20 |
| CALCRL | 2.4549E-12 | -0.526315939 | 0.667 | 0.702 | 6.0795E-08 |
| ATP5F1E | 3.19984E-31 | -0.525703543 | 0.91 | 0.96 | 7.9244E-27 |

| | | | | | |
|---|---|---|---|---|---|
| RPL35A | 2.09603E-44 | -0.523086091 | 0.836 | 0.957 | 5.1908E-40 |
| STMN1 | 4.2387E-18 | -0.522428273 | 0.14 | 0.299 | 1.0497E-13 |
| CAVIN1 | 3.66426E-25 | -0.5194249 | 0.394 | 0.556 | 9.0745E-21 |
| CFL1 | 1.03882E-30 | -0.518170641 | 0.868 | 0.918 | 2.5726E-26 |
| GAPDH | 7.91769E-31 | -0.516218683 | 0.819 | 0.923 | 1.9608E-26 |
| RPS12 | 8.93453E-40 | -0.515579484 | 0.936 | 0.989 | 2.2126E-35 |
| AHNAK | 1.00497E-20 | -0.514189771 | 0.251 | 0.428 | 2.4888E-16 |
| RPS25 | 5.42932E-45 | -0.511726442 | 0.811 | 0.947 | 1.3446E-40 |
| MTUS1 | 1.90122E-27 | -0.511545119 | 0.129 | 0.338 | 4.7084E-23 |
| RPL13 | 2.42717E-48 | -0.511447409 | 0.969 | 0.996 | 6.0109E-44 |
| CALD1 | 1.36128E-25 | -0.508804283 | 0.212 | 0.416 | 3.3712E-21 |
| RPL10 | 2.24702E-40 | -0.507419734 | 0.971 | 0.994 | 5.5647E-36 |
| GJA5 | 1.21288E-18 | -0.506386838 | 0.011 | 0.131 | 3.0037E-14 |
| RPL35 | 6.92288E-43 | -0.504349718 | 0.819 | 0.953 | 1.7145E-38 |
| GNAS | 9.33209E-32 | -0.502265833 | 0.85 | 0.86 | 2.3111E-27 |
| RPL18A | 8.11023E-45 | -0.501188638 | 0.859 | 0.973 | 2.0085E-40 |
| RPS27A | 4.14286E-48 | -0.500899501 | 0.916 | 0.985 | 1.026E-43 |
| RACK1 | 6.2846E-39 | -0.500587081 | 0.78 | 0.91 | 1.5564E-34 |
| VWA1 | 3.93482E-16 | -0.499863651 | 0.098 | 0.234 | 9.7446E-12 |
| RPS13 | 1.12632E-34 | -0.49907842 | 0.876 | 0.962 | 2.7893E-30 |
| CAVIN3 | 1.76459E-30 | -0.496432794 | 0.048 | 0.254 | 4.37E-26 |
| GUK1 | 2.89161E-26 | -0.493695196 | 0.529 | 0.674 | 7.1611E-22 |
| SOX18 | 1.05975E-14 | -0.493218914 | 0.361 | 0.487 | 2.6245E-10 |
| RPL19 | 9.24302E-42 | -0.48975596 | 0.853 | 0.98 | 2.289E-37 |
| MYL12B | 6.0927E-28 | -0.489614162 | 0.809 | 0.849 | 1.5089E-23 |
| RPL41 | 1.92579E-33 | -0.489277735 | 0.986 | 0.996 | 4.7692E-29 |
| ITGA6 | 1.21671E-19 | -0.488249514 | 0.126 | 0.291 | 3.0132E-15 |
| RPS3 | 7.50441E-43 | -0.486285775 | 0.833 | 0.973 | 1.8585E-38 |
| RPS21 | 1.26494E-41 | -0.484526104 | 0.727 | 0.924 | 3.1326E-37 |
| RPS15A | 1.52196E-39 | -0.480146096 | 0.851 | 0.974 | 3.7691E-35 |
| RPL12 | 3.66426E-36 | -0.479900363 | 0.825 | 0.962 | 9.0746E-32 |
| RPS27 | 5.62818E-38 | -0.464800402 | 0.949 | 0.993 | 1.3938E-33 |
| RPS7 | 6.51445E-34 | -0.461242247 | 0.865 | 0.968 | 1.6133E-29 |
| SELENOW | 7.21207E-16 | -0.459458853 | 0.633 | 0.696 | 1.7861E-11 |
| GNG11 | 1.08781E-20 | -0.459179035 | 0.724 | 0.841 | 2.694E-16 |

| | | | | | |
|---|---|---|---|---|---|
| MT-ND4L | 3.6783E-17 | -0.458777168 | 0.93 | 0.885 | 9.1093E-13 |
| CD320 | 3.48937E-17 | -0.458558347 | 0.105 | 0.254 | 8.6414E-13 |
| FABP5 | 5.56327E-12 | -0.456304518 | 0.068 | 0.178 | 1.3777E-07 |
| RPS4X | 4.5356E-41 | -0.455957525 | 0.719 | 0.952 | 1.1232E-36 |
| RPS14 | 1.33457E-47 | -0.453776791 | 0.86 | 0.979 | 3.3051E-43 |
| RPL22 | 4.42407E-36 | -0.453431305 | 0.772 | 0.903 | 1.0956E-31 |
| RPLP1 | 1.70534E-46 | -0.453136484 | 0.978 | 0.998 | 4.2233E-42 |
| RPS26 | 2.45296E-19 | -0.451512748 | 0.761 | 0.834 | 6.0748E-15 |
| FLNA | 4.26264E-28 | -0.450757084 | 0.096 | 0.309 | 1.0556E-23 |
| BCAM | 2.7889E-21 | -0.450451451 | 0.183 | 0.361 | 6.9067E-17 |
| SLCO2A1 | 4.50869E-24 | -0.442276058 | 0.034 | 0.199 | 1.1166E-19 |
| CD74 | 6.41579E-09 | -0.438584864 | 0.662 | 0.7 | 0.00015889 |
| RAB13 | 5.51371E-22 | -0.436370778 | 0.223 | 0.41 | 1.3655E-17 |
| RPL28 | 1.85582E-34 | -0.436342438 | 0.964 | 0.992 | 4.5959E-30 |
| PFDN5 | 1.60419E-29 | -0.432284286 | 0.667 | 0.828 | 3.9728E-25 |
| FAU | 2.00396E-38 | -0.43137494 | 0.874 | 0.962 | 4.9628E-34 |
| HEG1 | 1.16205E-10 | -0.430809348 | 0.327 | 0.419 | 2.8778E-06 |
| RPL5 | 1.63674E-30 | -0.430523351 | 0.743 | 0.907 | 4.0534E-26 |
| RPL18 | 1.98306E-38 | -0.429606573 | 0.89 | 0.971 | 4.9111E-34 |
| ACTG1 | 4.48633E-14 | -0.428684764 | 0.953 | 0.939 | 1.111E-09 |
| ANXA1 | 8.57285E-19 | -0.42785608 | 0.15 | 0.335 | 2.1231E-14 |
| ITGB1 | 3.7502E-15 | -0.42545589 | 0.718 | 0.773 | 9.2874E-11 |
| CPE | 9.14433E-07 | -0.42491112 | 0.074 | 0.143 | 0.02264593 |
| RPL15 | 1.39954E-33 | -0.423811849 | 0.885 | 0.976 | 3.466E-29 |
| MMP2 | 1.08476E-26 | -0.422698555 | 0.02 | 0.193 | 2.6864E-22 |
| RPS9 | 3.8741E-32 | -0.422494133 | 0.922 | 0.979 | 9.5942E-28 |
| OST4 | 2.0728E-16 | -0.421388782 | 0.645 | 0.701 | 5.1333E-12 |
| H3F3A | 5.17831E-25 | -0.419035778 | 0.845 | 0.9 | 1.2824E-20 |
| TSPO | 5.33695E-23 | -0.416482144 | 0.236 | 0.434 | 1.3217E-18 |
| RPL37A | 1.087E-31 | -0.415839356 | 0.834 | 0.944 | 2.6919E-27 |
| MT-CO2 | 2.75732E-22 | -0.414483802 | 0.995 | 0.997 | 6.8285E-18 |
| IGFBP2 | 2.16871E-28 | -0.414059026 | 0.016 | 0.195 | 5.3708E-24 |
| MYH9 | 3.3725E-15 | -0.409632781 | 0.397 | 0.515 | 8.352E-11 |
| TMA7 | 9.60937E-22 | -0.40915838 | 0.671 | 0.772 | 2.3798E-17 |
| ADAM15 | 1.21887E-14 | -0.407548742 | 0.202 | 0.341 | 3.0185E-10 |

| | | | | | |
|---|---|---|---|---|---|
| MT-CYB | 2.38776E-20 | -0.407154573 | 0.974 | 0.988 | 5.9133E-16 |
| THSD7A | 1.28499E-17 | -0.406548665 | 0.11 | 0.261 | 3.1823E-13 |
| RPL14 | 5.25741E-36 | -0.406336951 | 0.769 | 0.937 | 1.302E-31 |
| RPS5 | 2.11464E-44 | -0.40572031 | 0.598 | 0.871 | 5.2369E-40 |
| GNG5 | 4.10919E-17 | -0.40480127 | 0.442 | 0.574 | 1.0176E-12 |
| EEF1A1 | 1.15586E-34 | -0.404345309 | 0.995 | 0.999 | 2.8625E-30 |
| CCDC85B | 3.39141E-16 | -0.403630523 | 0.64 | 0.706 | 8.3988E-12 |
| HES4 | 3.56944E-15 | -0.398534269 | 0.104 | 0.239 | 8.8397E-11 |
| MT-ND3 | 7.02229E-21 | -0.397111363 | 0.995 | 0.995 | 1.7391E-16 |
| MT-ND4 | 5.52329E-21 | -0.394222015 | 0.995 | 0.994 | 1.3678E-16 |
| LTBP4 | 2.11922E-17 | -0.394132894 | 0.04 | 0.17 | 5.2483E-13 |
| SYNPO | 9.19528E-29 | -0.392419834 | 0.064 | 0.267 | 2.2772E-24 |
| RPL24 | 4.51765E-21 | -0.383019558 | 0.909 | 0.95 | 1.1188E-16 |
| EEF1D | 4.04778E-30 | -0.381775444 | 0.67 | 0.828 | 1.0024E-25 |
| GJA1 | 3.42196E-22 | -0.380452543 | 0.028 | 0.18 | 8.4745E-18 |
| RPL6 | 2.47861E-29 | -0.379615923 | 0.797 | 0.943 | 6.1383E-25 |
| PFN1 | 1.35346E-12 | -0.377300748 | 0.814 | 0.85 | 3.3519E-08 |
| MDK | 2.38748E-20 | -0.376480696 | 0.105 | 0.276 | 5.9126E-16 |
| RPLP2 | 1.84619E-30 | -0.371481146 | 0.896 | 0.97 | 4.5721E-26 |
| RPL23 | 2.32933E-19 | -0.371022085 | 0.792 | 0.851 | 5.7686E-15 |
| RPL10A | 1.24474E-37 | -0.370712959 | 0.588 | 0.855 | 3.0826E-33 |
| COX6C | 2.164E-15 | -0.369199737 | 0.54 | 0.654 | 5.3591E-11 |
| SH3BGRL3 | 3.35919E-25 | -0.367543168 | 0.608 | 0.757 | 8.319E-21 |
| PPIA | 1.66045E-15 | -0.366586394 | 0.921 | 0.914 | 4.1121E-11 |
| TUBB | 1.03992E-11 | -0.364146536 | 0.344 | 0.462 | 2.5754E-07 |
| RPL39 | 9.46194E-17 | -0.364058537 | 0.966 | 0.979 | 2.3432E-12 |
| HINT1 | 9.9809E-17 | -0.363236455 | 0.544 | 0.669 | 2.4718E-12 |
| SNHG7 | 1.49076E-11 | -0.361741342 | 0.251 | 0.371 | 3.6919E-07 |
| MT-CO1 | 1.538E-15 | -0.357266431 | 0.997 | 0.996 | 3.8088E-11 |
| LMO2 | 2.95878E-12 | -0.35308872 | 0.242 | 0.365 | 7.3274E-08 |
| GSTP1 | 8.99752E-15 | -0.348597527 | 0.429 | 0.555 | 2.2282E-10 |
| MYL12A | 2.83792E-09 | -0.348210143 | 0.91 | 0.882 | 7.0281E-05 |
| MT-CO3 | 7.93693E-18 | -0.34733053 | 0.981 | 0.99 | 1.9656E-13 |
| ARL15 | 8.95027E-09 | -0.345486978 | 0.188 | 0.283 | 0.00022165 |
| SEMA3G | 6.46305E-19 | -0.336203938 | 0.006 | 0.124 | 1.6006E-14 |

| | | | | | |
|---|---|---|---|---|---|
| LRRFIP1 | 4.02419E-15 | -0.335672148 | 0.178 | 0.329 | 9.9659E-11 |
| ENTPD1 | 3.21427E-26 | -0.334494288 | 0.037 | 0.217 | 7.9601E-22 |
| RPS11 | 2.93052E-25 | -0.333699446 | 0.786 | 0.909 | 7.2574E-21 |
| UBA52 | 4.02734E-29 | -0.331016743 | 0.786 | 0.912 | 9.9737E-25 |
| ECSCR | 5.29591E-19 | -0.330776199 | 0.203 | 0.378 | 1.3115E-14 |
| ELOB | 5.33884E-11 | -0.329464374 | 0.553 | 0.633 | 1.3222E-06 |
| NDUFB2 | 6.68701E-12 | -0.328890518 | 0.42 | 0.529 | 1.656E-07 |
| EMCN | 2.90542E-09 | -0.32736034 | 0.358 | 0.451 | 7.1953E-05 |
| LAMB1 | 1.87985E-22 | -0.327287145 | 0.031 | 0.186 | 4.6554E-18 |
| TUBA1A | 1.33061E-09 | -0.32239809 | 0.217 | 0.328 | 3.2953E-05 |
| HSPB1 | 2.26576E-09 | -0.321675046 | 0.581 | 0.644 | 5.6111E-05 |
| LAMA4 | 1.05275E-15 | -0.321356604 | 0.109 | 0.251 | 2.6071E-11 |
| MGST3 | 4.54248E-11 | -0.318728782 | 0.336 | 0.453 | 1.1249E-06 |
| BTF3 | 8.46381E-16 | -0.318688121 | 0.701 | 0.781 | 2.0961E-11 |
| POLR2L | 1.33884E-08 | -0.317607281 | 0.656 | 0.689 | 0.00033156 |
| MCTP1 | 3.77167E-17 | -0.31712964 | 0.053 | 0.187 | 9.3405E-13 |
| UBL5 | 1.47724E-10 | -0.317019566 | 0.535 | 0.613 | 3.6584E-06 |
| MYL6 | 1.05106E-12 | -0.316151046 | 0.989 | 0.975 | 2.6029E-08 |
| EDF1 | 6.41535E-12 | -0.314928574 | 0.509 | 0.61 | 1.5888E-07 |
| THBS1 | 2.74547E-07 | -0.314542673 | 0.068 | 0.141 | 0.00679916 |
| DST | 6.94801E-13 | -0.314529771 | 0.143 | 0.267 | 1.7207E-08 |
| DSTN | 2.1014E-09 | -0.314377097 | 0.516 | 0.59 | 5.2041E-05 |
| JAG2 | 4.81863E-22 | -0.313707139 | 0.028 | 0.18 | 1.1933E-17 |
| APLP2 | 3.32275E-10 | -0.313505427 | 0.291 | 0.399 | 8.2288E-06 |
| NAA38 | 1.23891E-10 | -0.311843278 | 0.38 | 0.493 | 3.0682E-06 |
| RPS29 | 1.90491E-18 | -0.309395279 | 0.888 | 0.958 | 4.7175E-14 |
| RPL38 | 2.61649E-16 | -0.308509609 | 0.82 | 0.91 | 6.4797E-12 |
| LMCD1 | 1.42782E-21 | -0.305225422 | 0.022 | 0.167 | 3.536E-17 |
| C4orf48 | 1.03117E-09 | -0.304880854 | 0.248 | 0.353 | 2.5537E-05 |
| TOMM7 | 1.51931E-13 | -0.303203424 | 0.54 | 0.647 | 3.7626E-09 |
| SNX3 | 6.14202E-09 | -0.301828954 | 0.443 | 0.529 | 0.00015211 |
| RPL27 | 2.53164E-20 | -0.301696857 | 0.719 | 0.878 | 6.2696E-16 |
| SULF2 | 6.2643E-16 | -0.299565962 | 0.143 | 0.291 | 1.5514E-11 |
| RPL27A | 5.692E-23 | -0.2982123 | 0.788 | 0.907 | 1.4096E-18 |
| TMTC1 | 3.61006E-19 | -0.297010708 | 0.028 | 0.161 | 8.9403E-15 |

| | | | | | |
|---|---|---|---|---|---|
| ELN | 1.36906E-17 | -0.296974163 | 0.008 | 0.119 | 3.3905E-13 |
| MT-ND5 | 7.47119E-11 | -0.296261269 | 0.899 | 0.919 | 1.8502E-06 |
| VEGFC | 1.13353E-19 | -0.295685422 | 0.016 | 0.145 | 2.8072E-15 |
| SWAP70 | 2.92524E-11 | -0.294216775 | 0.214 | 0.335 | 7.2444E-07 |
| FIS1 | 2.21445E-11 | -0.292722844 | 0.369 | 0.487 | 5.4841E-07 |
| MECOM | 1.52132E-11 | -0.292555883 | 0.119 | 0.234 | 3.7675E-07 |
| NUCKS1 | 2.17516E-08 | -0.291479199 | 0.329 | 0.417 | 0.00053868 |
| COMMD6 | 1.02572E-09 | -0.291205777 | 0.408 | 0.503 | 2.5402E-05 |
| GABARAPL2 | 8.25996E-09 | -0.291187313 | 0.353 | 0.442 | 0.00020456 |
| PLEC | 7.44077E-12 | -0.289963277 | 0.174 | 0.299 | 1.8427E-07 |
| CYYR1 | 3.28701E-07 | -0.287883724 | 0.363 | 0.427 | 0.00814028 |
| MT-ATP6 | 1.57896E-13 | -0.284692866 | 0.994 | 0.994 | 3.9103E-09 |
| NEDD8 | 2.05499E-10 | -0.280289292 | 0.296 | 0.414 | 5.0892E-06 |
| ELK3 | 2.80465E-08 | -0.280209717 | 0.412 | 0.494 | 0.00069457 |
| HMGB1 | 5.11046E-09 | -0.279756769 | 0.867 | 0.879 | 0.00012656 |
| FBN1 | 8.86559E-15 | -0.278482814 | 0.079 | 0.207 | 2.1956E-10 |
| COX8A | 1.09956E-07 | -0.278341089 | 0.381 | 0.462 | 0.00272305 |
| MZT2B | 2.0945E-09 | -0.273910782 | 0.322 | 0.431 | 5.187E-05 |
| JAM2 | 1.01946E-08 | -0.271020199 | 0.219 | 0.316 | 0.00025247 |
| MPZL2 | 3.22248E-23 | -0.270941108 | 0.003 | 0.143 | 7.9805E-19 |
| SNCG | 5.24256E-10 | -0.269442532 | 0.144 | 0.25 | 1.2983E-05 |
| BLOC1S1 | 1.13475E-13 | -0.26883098 | 0.174 | 0.318 | 2.8102E-09 |
| TPM4 | 3.09764E-07 | -0.268673718 | 0.695 | 0.725 | 0.00767131 |
| LIMS1 | 8.70338E-10 | -0.267095723 | 0.16 | 0.268 | 2.1554E-05 |
| ZEB1 | 1.54009E-11 | -0.266556659 | 0.2 | 0.326 | 3.814E-07 |
| IFITM1 | 1.12947E-06 | -0.265078889 | 0.223 | 0.304 | 0.02797141 |
| MT-ND1 | 3.88507E-08 | -0.264806625 | 0.983 | 0.974 | 0.00096214 |
| MAP4 | 7.5788E-10 | -0.263152688 | 0.209 | 0.322 | 1.8769E-05 |
| EID1 | 3.42045E-08 | -0.262621131 | 0.442 | 0.516 | 0.00084708 |
| ACVRL1 | 6.03708E-14 | -0.262106685 | 0.133 | 0.267 | 1.4951E-09 |
| SUMO2 | 1.59242E-06 | -0.261822896 | 0.633 | 0.651 | 0.03943621 |
| UQCRB | 1.92208E-10 | -0.261381918 | 0.519 | 0.605 | 4.76E-06 |
| ABLIM1 | 2.9541E-16 | -0.260584581 | 0.074 | 0.211 | 7.3158E-12 |
| MAP4K4 | 1.00338E-09 | -0.260396424 | 0.242 | 0.357 | 2.4849E-05 |
| RPL31 | 1.11358E-07 | -0.259193958 | 0.481 | 0.548 | 0.00275778 |

| | | | | | |
|---|---|---|---|---|---|
| UQCRQ | 3.63324E-07 | -0.255846991 | 0.33 | 0.415 | 0.00899771 |
| IFITM2 | 9.58322E-09 | -0.254142459 | 0.797 | 0.8 | 0.00023733 |
| ATP5PF | 6.34025E-09 | -0.253713259 | 0.426 | 0.515 | 0.00015702 |
| C19orf53 | 1.78603E-09 | -0.252834265 | 0.25 | 0.364 | 4.4231E-05 |
| RPL36AL | 1.27793E-10 | -0.252620901 | 0.592 | 0.674 | 3.1648E-06 |
| RPL21 | 8.86257E-28 | -0.25171933 | 0.812 | 0.945 | 2.1948E-23 |
| PPDPF | 1.86916E-08 | -0.251511334 | 0.304 | 0.402 | 0.0004629 |
| ABCG2 | 6.45289E-13 | -0.250561411 | 0.056 | 0.165 | 1.5981E-08 |
| ADAMTS6 | 5.79697E-09 | -0.250417237 | 0.05 | 0.129 | 0.00014356 |
| DNAJB14 | 5.31208E-21 | 0.250951797 | 0.375 | 0.199 | 1.3155E-16 |
| LINC02388 | 1.3122E-46 | 0.250977344 | 0.228 | 0.051 | 3.2497E-42 |
| WDFY2 | 1.74278E-39 | 0.251620095 | 0.222 | 0.057 | 4.316E-35 |
| IGF1R | 2.23891E-22 | 0.251687495 | 0.318 | 0.152 | 5.5447E-18 |
| MGAT1 | 1.06121E-16 | 0.252528888 | 0.502 | 0.326 | 2.6281E-12 |
| TAF1D | 1.23507E-19 | 0.25264979 | 0.375 | 0.204 | 3.0587E-15 |
| RREB1 | 3.07252E-29 | 0.252663568 | 0.262 | 0.097 | 7.6091E-25 |
| ENOSF1 | 2.40011E-27 | 0.252741681 | 0.281 | 0.113 | 5.9439E-23 |
| NRP2 | 1.66366E-17 | 0.252833101 | 0.549 | 0.353 | 4.1201E-13 |
| SLC30A7 | 1.11717E-22 | 0.253193399 | 0.335 | 0.166 | 2.7667E-18 |
| SF1 | 4.76383E-15 | 0.25320776 | 0.468 | 0.308 | 1.1798E-10 |
| CYTH1 | 1.06544E-26 | 0.253824514 | 0.374 | 0.175 | 2.6386E-22 |
| IRF1 | 3.44537E-19 | 0.253865514 | 0.371 | 0.198 | 8.5325E-15 |
| ERV3-1 | 6.95609E-42 | 0.254149951 | 0.186 | 0.038 | 1.7227E-37 |
| CHSY1 | 4.1941E-20 | 0.254425263 | 0.448 | 0.26 | 1.0387E-15 |
| HELZ | 1.79888E-21 | 0.254753798 | 0.35 | 0.18 | 4.4549E-17 |
| ATRNL1 | 1.8935E-41 | 0.255157465 | 0.288 | 0.087 | 4.6893E-37 |
| TCF7L2 | 1.03559E-20 | 0.255171268 | 0.363 | 0.192 | 2.5646E-16 |
| ANKRD12 | 1.73061E-19 | 0.255729955 | 0.485 | 0.289 | 4.2858E-15 |
| CDK17 | 5.6344E-23 | 0.255837255 | 0.505 | 0.293 | 1.3954E-18 |
| ZNF844 | 5.99513E-41 | 0.255868394 | 0.225 | 0.057 | 1.4847E-36 |
| FARP2 | 9.27291E-42 | 0.255976181 | 0.233 | 0.059 | 2.2964E-37 |
| ZEB2 | 1.27848E-20 | 0.255978318 | 0.322 | 0.159 | 3.1661E-16 |
| PROS1 | 1.20977E-17 | 0.255998683 | 0.512 | 0.332 | 2.996E-13 |
| MEPCE | 9.93868E-41 | 0.256404496 | 0.264 | 0.076 | 2.4613E-36 |
| KIAA1551 | 2.25532E-17 | 0.257463886 | 0.35 | 0.192 | 5.5853E-13 |

| | | | | | |
|---|---|---|---|---|---|
| MTRF1L | 2.05268E-21 | 0.257471619 | 0.374 | 0.193 | 5.0835E-17 |
| B4GALT3 | 1.39671E-33 | 0.257805121 | 0.273 | 0.095 | 3.4589E-29 |
| STOM | 5.84576E-10 | 0.257865751 | 0.764 | 0.702 | 1.4477E-05 |
| CABLES1 | 2.04356E-38 | 0.258582553 | 0.285 | 0.09 | 5.0609E-34 |
| IER5 | 7.97773E-19 | 0.258637042 | 0.322 | 0.167 | 1.9757E-14 |
| LAMP2 | 2.6439E-20 | 0.25870116 | 0.518 | 0.319 | 6.5476E-16 |
| AL118516.1 | 6.27865E-28 | 0.25901191 | 0.22 | 0.075 | 1.5549E-23 |
| BCL2L11 | 2.73189E-18 | 0.259016469 | 0.336 | 0.179 | 6.7655E-14 |
| FAM43A | 8.73969E-22 | 0.259115976 | 0.371 | 0.194 | 2.1644E-17 |
| HNRNPDL | 1.61049E-15 | 0.259186484 | 0.732 | 0.567 | 3.9884E-11 |
| LTK | 1.52799E-53 | 0.259308482 | 0.216 | 0.038 | 3.7841E-49 |
| Mar-07 | 4.61615E-24 | 0.259592087 | 0.411 | 0.213 | 1.1432E-19 |
| TFPI2 | 9.00947E-15 | 0.259652598 | 0.425 | 0.258 | 2.2312E-10 |
| ELMSAN1 | 1.91226E-27 | 0.260415896 | 0.394 | 0.188 | 4.7357E-23 |
| SETD5 | 7.42442E-27 | 0.260534587 | 0.377 | 0.181 | 1.8387E-22 |
| PTPN1 | 1.13244E-20 | 0.260751899 | 0.329 | 0.163 | 2.8045E-16 |
| ZSWIM6 | 4.1721E-29 | 0.26088552 | 0.318 | 0.134 | 1.0332E-24 |
| MORF4L2 | 6.08475E-18 | 0.262089704 | 0.437 | 0.266 | 1.5069E-13 |
| F2RL3 | 1.24979E-22 | 0.262374141 | 0.344 | 0.167 | 3.0951E-18 |
| LRRFIP2 | 7.06532E-26 | 0.262493841 | 0.433 | 0.225 | 1.7497E-21 |
| TIPARP | 2.8337E-25 | 0.262548388 | 0.302 | 0.13 | 7.0176E-21 |
| SNX13 | 3.21804E-22 | 0.26307003 | 0.357 | 0.183 | 7.9695E-18 |
| TLE4 | 1.12239E-14 | 0.263824226 | 0.273 | 0.148 | 2.7796E-10 |
| TSPAN14 | 2.35913E-17 | 0.264288776 | 0.515 | 0.335 | 5.8424E-13 |
| SH3TC1 | 3.47774E-28 | 0.264422978 | 0.343 | 0.151 | 8.6126E-24 |
| CEACAM1 | 1.03083E-41 | 0.264441622 | 0.278 | 0.082 | 2.5529E-37 |
| KALRN | 2.99388E-33 | 0.26494719 | 0.374 | 0.158 | 7.4143E-29 |
| NISCH | 1.60717E-28 | 0.266276015 | 0.326 | 0.14 | 3.9802E-24 |
| APOL6 | 2.90315E-26 | 0.266367628 | 0.358 | 0.166 | 7.1897E-22 |
| SNX18 | 1.35629E-27 | 0.266630288 | 0.302 | 0.126 | 3.3589E-23 |
| MKLN1 | 4.70897E-21 | 0.267308034 | 0.355 | 0.185 | 1.1662E-16 |
| SERPING1 | 8.69035E-20 | 0.26763635 | 0.518 | 0.325 | 2.1522E-15 |
| SERPINH1 | 1.04201E-19 | 0.268188658 | 0.6 | 0.396 | 2.5805E-15 |
| HINFP | 1.71062E-52 | 0.268515362 | 0.223 | 0.043 | 4.2363E-48 |
| SELENOS | 8.38349E-18 | 0.268611562 | 0.481 | 0.306 | 2.0762E-13 |

| | | | | | |
|---|---|---|---|---|---|
| PLAUR | 3.76876E-26 | 0.269386256 | 0.268 | 0.107 | 9.3333E-22 |
| TPP1 | 1.08762E-27 | 0.26983915 | 0.474 | 0.245 | 2.6935E-23 |
| SIK2 | 1.04416E-26 | 0.270011672 | 0.315 | 0.138 | 2.5859E-22 |
| CCNH | 6.54926E-19 | 0.271851855 | 0.29 | 0.144 | 1.6219E-14 |
| ORM2 | 3.16603E-20 | 0.27262803 | 0.101 | 0.024 | 7.8407E-16 |
| SEMA4C | 1.05842E-25 | 0.2726406 | 0.316 | 0.141 | 2.6212E-21 |
| MARCKSL1 | 2.11329E-13 | 0.273531715 | 0.774 | 0.674 | 5.2336E-09 |
| LAPTM4B | 3.37007E-21 | 0.274365314 | 0.437 | 0.248 | 8.346E-17 |
| SFMBT2 | 1.28307E-38 | 0.274584649 | 0.27 | 0.084 | 3.1775E-34 |
| PDK1 | 3.83153E-44 | 0.275586148 | 0.231 | 0.056 | 9.4888E-40 |
| SLC25A37 | 1.86968E-24 | 0.275597077 | 0.47 | 0.256 | 4.6303E-20 |
| MAFB | 1.14172E-19 | 0.276007534 | 0.197 | 0.077 | 2.8275E-15 |
| SERHL2 | 2.24381E-51 | 0.27610278 | 0.229 | 0.046 | 5.5568E-47 |
| PITPNB | 2.8211E-21 | 0.276185537 | 0.434 | 0.249 | 6.9865E-17 |
| C2CD4B | 1.59941E-24 | 0.276185537 | 0.363 | 0.173 | 3.9609E-20 |
| MAT2A | 2.02632E-23 | 0.276818501 | 0.377 | 0.19 | 5.0182E-19 |
| AC016831.5 | 3.32539E-45 | 0.277257906 | 0.208 | 0.044 | 8.2353E-41 |
| SNX5 | 1.32417E-28 | 0.277319115 | 0.456 | 0.229 | 3.2793E-24 |
| INMT | 1.71142E-20 | 0.27850141 | 0.429 | 0.233 | 4.2383E-16 |
| SON | 3.8639E-17 | 0.278698726 | 0.726 | 0.541 | 9.5689E-13 |
| TCP1 | 1.12057E-19 | 0.27886908 | 0.426 | 0.248 | 2.7751E-15 |
| SYAP1 | 8.92017E-25 | 0.27886908 | 0.437 | 0.229 | 2.2091E-20 |
| SHANK3 | 1.81783E-20 | 0.280358681 | 0.51 | 0.314 | 4.5019E-16 |
| RHOU | 1.44067E-23 | 0.280512173 | 0.346 | 0.166 | 3.5678E-19 |
| SNX8 | 1.1799E-37 | 0.280568371 | 0.394 | 0.156 | 2.922E-33 |
| TMEM123 | 1.31868E-15 | 0.280715751 | 0.608 | 0.443 | 3.2657E-11 |
| MAP3K6 | 5.96516E-34 | 0.280814441 | 0.353 | 0.144 | 1.4773E-29 |
| ZNF347 | 2.46897E-34 | 0.281581646 | 0.304 | 0.113 | 6.1144E-30 |
| GLO1 | 5.19055E-24 | 0.284213399 | 0.526 | 0.304 | 1.2854E-19 |
| GCLM | 2.88082E-32 | 0.284243459 | 0.347 | 0.143 | 7.1343E-28 |
| GPX3 | 1.67765E-18 | 0.284577947 | 0.414 | 0.237 | 4.1547E-14 |
| TGFB3 | 1.53987E-49 | 0.284923223 | 0.243 | 0.055 | 3.8135E-45 |
| P4HB | 1.87314E-18 | 0.285469802 | 0.625 | 0.436 | 4.6388E-14 |
| EWSR1 | 1.24013E-21 | 0.287393545 | 0.451 | 0.261 | 3.0712E-17 |
| PLXNA2 | 5.14648E-29 | 0.287596547 | 0.355 | 0.159 | 1.2745E-24 |

| | | | | | |
|---|---|---|---|---|---|
| GATA6 | 1.2906E-39 | 0.288388367 | 0.285 | 0.09 | 3.1962E-35 |
| ROBO4 | 1.76791E-19 | 0.28917798 | 0.594 | 0.395 | 4.3782E-15 |
| UGP2 | 9.43434E-21 | 0.289277769 | 0.302 | 0.148 | 2.3364E-16 |
| PDIA3 | 5.19373E-14 | 0.289316901 | 0.702 | 0.585 | 1.2862E-09 |
| TRIM35 | 1.94284E-42 | 0.28952067 | 0.358 | 0.125 | 4.8115E-38 |
| SNHG8 | 4.49003E-15 | 0.28969611 | 0.377 | 0.23 | 1.112E-10 |
| CALU | 4.66019E-18 | 0.289701904 | 0.606 | 0.424 | 1.1541E-13 |
| TMEM165 | 2.12743E-21 | 0.290887584 | 0.488 | 0.296 | 5.2686E-17 |
| CTSB | 1.43886E-17 | 0.292381105 | 0.617 | 0.432 | 3.5633E-13 |
| VTN | 3.58491E-15 | 0.292494519 | 0.126 | 0.046 | 8.878E-11 |
| TSPAN7 | 2.06893E-21 | 0.292639796 | 0.643 | 0.425 | 5.1237E-17 |
| MT2A | 1.59709E-16 | 0.292991994 | 0.94 | 0.824 | 3.9552E-12 |
| DSE | 2.40065E-38 | 0.293518883 | 0.336 | 0.123 | 5.9452E-34 |
| CHST15 | 4.52646E-37 | 0.293591731 | 0.338 | 0.126 | 1.121E-32 |
| TMEM50B | 7.21108E-30 | 0.293964842 | 0.462 | 0.235 | 1.7858E-25 |
| PXN | 6.56128E-24 | 0.294347474 | 0.467 | 0.265 | 1.6249E-19 |
| TUBA1C | 1.2873E-25 | 0.294666657 | 0.384 | 0.19 | 3.188E-21 |
| SETX | 2.66131E-21 | 0.294987458 | 0.505 | 0.307 | 6.5907E-17 |
| GNAI3 | 1.03363E-20 | 0.295097325 | 0.457 | 0.272 | 2.5598E-16 |
| NEDD9 | 3.65947E-14 | 0.295205707 | 0.591 | 0.436 | 9.0627E-10 |
| ASAH1 | 5.10194E-24 | 0.295575614 | 0.544 | 0.323 | 1.2635E-19 |
| ERVK3-1 | 1.217E-36 | 0.295816015 | 0.276 | 0.092 | 3.0139E-32 |
| TOB2 | 5.14714E-27 | 0.296058571 | 0.361 | 0.171 | 1.2747E-22 |
| NKTR | 2.2907E-22 | 0.296139908 | 0.509 | 0.3 | 5.6729E-18 |
| PARP14 | 1.98341E-17 | 0.296397929 | 0.55 | 0.375 | 4.9119E-13 |
| DCXR | 1.92708E-07 | 0.296847998 | 0.18 | 0.111 | 0.00477241 |
| ARL5B | 3.54155E-28 | 0.297317188 | 0.388 | 0.185 | 8.7707E-24 |
| CNN3 | 7.73439E-13 | 0.297698258 | 0.718 | 0.62 | 1.9154E-08 |
| MEG8 | 2.57325E-56 | 0.298085573 | 0.197 | 0.029 | 6.3726E-52 |
| NEU1 | 8.39984E-33 | 0.298651345 | 0.307 | 0.118 | 2.0802E-28 |
| PABPC4 | 3.94045E-27 | 0.299036791 | 0.374 | 0.18 | 9.7585E-23 |
| NR5A2 | 7.64429E-34 | 0.299052496 | 0.353 | 0.142 | 1.8931E-29 |
| FAM53C | 7.94E-47 | 0.300940501 | 0.259 | 0.066 | 1.9663E-42 |
| MAP7D3 | 5.99364E-31 | 0.301250049 | 0.316 | 0.125 | 1.4843E-26 |
| GPM6A | 2.45174E-26 | 0.301516216 | 0.451 | 0.232 | 6.0717E-22 |

| | | | | | |
|---|---|---|---|---|---|
| FGB | 3.37359E-14 | 0.301881221 | 0.116 | 0.042 | 8.3547E-10 |
| SERPINB9 | 7.78654E-22 | 0.30225963 | 0.419 | 0.231 | 1.9283E-17 |
| RNF149 | 4.40033E-41 | 0.302312842 | 0.31 | 0.101 | 1.0897E-36 |
| EPB41L2 | 3.69115E-23 | 0.303567177 | 0.389 | 0.206 | 9.1411E-19 |
| ZFP36L1 | 5.51547E-15 | 0.304621133 | 0.645 | 0.477 | 1.3659E-10 |
| EIF4A3 | 7.57846E-27 | 0.305291518 | 0.381 | 0.182 | 1.8768E-22 |
| CCL23 | 6.82707E-28 | 0.306046404 | 0.344 | 0.149 | 1.6907E-23 |
| DAPK1 | 1.33289E-38 | 0.306275169 | 0.366 | 0.14 | 3.3009E-34 |
| PLXDC2 | 4.78388E-21 | 0.307894534 | 0.408 | 0.23 | 1.1847E-16 |
| SHC1 | 1.96803E-25 | 0.308371043 | 0.527 | 0.312 | 4.8738E-21 |
| BCL2L13 | 5.99853E-49 | 0.308424354 | 0.326 | 0.097 | 1.4855E-44 |
| PLEKHO2 | 4.04099E-37 | 0.309928641 | 0.402 | 0.169 | 1.0008E-32 |
| PMAIP1 | 4.79399E-37 | 0.310341402 | 0.198 | 0.049 | 1.1872E-32 |
| ARL4D | 4.566E-33 | 0.310640347 | 0.285 | 0.103 | 1.1308E-28 |
| CNST | 5.28891E-37 | 0.311008632 | 0.419 | 0.181 | 1.3098E-32 |
| STAT3 | 6.99151E-17 | 0.311193113 | 0.555 | 0.399 | 1.7314E-12 |
| PPP1R15B | 2.09771E-34 | 0.31154379 | 0.406 | 0.177 | 5.195E-30 |
| ECM1 | 6.70207E-39 | 0.312620237 | 0.36 | 0.132 | 1.6598E-34 |
| PELI1 | 3.17818E-22 | 0.312752263 | 0.501 | 0.304 | 7.8708E-18 |
| DNMBP | 1.61342E-55 | 0.313912129 | 0.291 | 0.071 | 3.9956E-51 |
| TOB1 | 5.10038E-30 | 0.314452234 | 0.296 | 0.118 | 1.2631E-25 |
| IL7R | 2.67485E-17 | 0.315517404 | 0.105 | 0.03 | 6.6243E-13 |
| DNAJB6 | 1.11729E-21 | 0.316802719 | 0.557 | 0.351 | 2.767E-17 |
| NUTM2B-AS1 | 8.13473E-37 | 0.317970532 | 0.324 | 0.122 | 2.0146E-32 |
| GPR182 | 2.62489E-48 | 0.318559779 | 0.324 | 0.096 | 6.5005E-44 |
| SEC14L1 | 8.08653E-15 | 0.319644146 | 0.777 | 0.686 | 2.0026E-10 |
| UBB | 2.56469E-17 | 0.320571874 | 0.96 | 0.897 | 6.3515E-13 |
| C16orf72 | 5.34972E-30 | 0.320744366 | 0.381 | 0.177 | 1.3249E-25 |
| NR2F1-AS1 | 7.93977E-53 | 0.321191036 | 0.321 | 0.09 | 1.9663E-48 |
| MAN1A1 | 3.51978E-31 | 0.321314277 | 0.487 | 0.248 | 8.7167E-27 |
| SASH1 | 1.29822E-22 | 0.321457076 | 0.584 | 0.378 | 3.215E-18 |
| DNTTIP2 | 8.72687E-23 | 0.322517167 | 0.394 | 0.213 | 2.1612E-18 |
| DHCR24 | 1.15432E-43 | 0.32312024 | 0.335 | 0.11 | 2.8587E-39 |
| RPGR | 4.18903E-31 | 0.32316711 | 0.42 | 0.203 | 1.0374E-26 |
| MYSM1 | 3.21312E-41 | 0.323465454 | 0.381 | 0.148 | 7.9573E-37 |

| | | | | | |
|---|---|---|---|---|---|
| ST3GAL4 | 2.48436E-32 | 0.323802366 | 0.35 | 0.148 | 6.1525E-28 |
| DLC1 | 1.26258E-18 | 0.324157366 | 0.6 | 0.412 | 3.1268E-14 |
| SERPINB1 | 7.83032E-22 | 0.324440671 | 0.53 | 0.324 | 1.9392E-17 |
| CXCL8 | 2.21386E-26 | 0.324573177 | 0.236 | 0.086 | 5.4826E-22 |
| CPD | 2.51313E-23 | 0.324605845 | 0.447 | 0.254 | 6.2238E-19 |
| CDK11B | 3.33039E-38 | 0.324742321 | 0.291 | 0.098 | 8.2477E-34 |
| SIRT1 | 9.29999E-47 | 0.326207959 | 0.281 | 0.079 | 2.3031E-42 |
| FGG | 2.05932E-17 | 0.326835416 | 0.124 | 0.04 | 5.0999E-13 |
| NRP1 | 2.07091E-17 | 0.326888444 | 0.712 | 0.559 | 5.1286E-13 |
| ART4 | 3.40099E-44 | 0.327448244 | 0.383 | 0.137 | 8.4226E-40 |
| RBM5 | 6.61204E-30 | 0.328081768 | 0.433 | 0.218 | 1.6375E-25 |
| ICAM4 | 6.9279E-78 | 0.328314192 | 0.229 | 0.026 | 1.7157E-73 |
| TINAGL1 | 7.05229E-17 | 0.328506277 | 0.69 | 0.532 | 1.7465E-12 |
| UBR1 | 3.13541E-41 | 0.329356653 | 0.34 | 0.122 | 7.7648E-37 |
| NASP | 6.00848E-22 | 0.329487188 | 0.493 | 0.302 | 1.488E-17 |
| SLC9A9 | 1.51578E-37 | 0.330121302 | 0.409 | 0.173 | 3.7538E-33 |
| MIDN | 2.09295E-23 | 0.33013889 | 0.637 | 0.423 | 5.1832E-19 |
| FARP1 | 9.43333E-44 | 0.330184788 | 0.307 | 0.098 | 2.3362E-39 |
| NRCAM | 4.58441E-51 | 0.331326146 | 0.299 | 0.081 | 1.1353E-46 |
| ZNF331 | 1.0891E-41 | 0.331563467 | 0.243 | 0.065 | 2.6972E-37 |
| CAVIN2 | 7.67715E-10 | 0.333103006 | 0.76 | 0.693 | 1.9012E-05 |
| B4GALT5 | 1.81022E-24 | 0.333125331 | 0.453 | 0.253 | 4.483E-20 |
| NDUFS2 | 7.39053E-28 | 0.333741732 | 0.465 | 0.249 | 1.8303E-23 |
| MID1 | 2.5806E-43 | 0.333832823 | 0.408 | 0.159 | 6.3909E-39 |
| HSP90B1 | 6.18579E-12 | 0.335758369 | 0.795 | 0.757 | 1.5319E-07 |
| FOSL2 | 1.23364E-19 | 0.336045105 | 0.569 | 0.382 | 3.0551E-15 |
| SLC16A6 | 3.14397E-44 | 0.336229717 | 0.211 | 0.046 | 7.786E-40 |
| TNFRSF1A | 2.76872E-24 | 0.336840362 | 0.6 | 0.393 | 6.8567E-20 |
| UBE2S | 1.07412E-26 | 0.338618083 | 0.346 | 0.16 | 2.6601E-22 |
| ARHGEF7 | 2.77806E-34 | 0.339352098 | 0.487 | 0.245 | 6.8799E-30 |
| SDCBP | 2.04706E-19 | 0.339642696 | 0.743 | 0.603 | 5.0695E-15 |
| CNOT6L | 1.44744E-17 | 0.340176499 | 0.471 | 0.308 | 3.5846E-13 |
| NFKBIA | 7.01434E-21 | 0.340457206 | 0.876 | 0.722 | 1.7371E-16 |
| CCNT1 | 3.53633E-32 | 0.341490126 | 0.338 | 0.139 | 8.7577E-28 |
| GSTA1 | 5.49853E-17 | 0.341658041 | 0.121 | 0.039 | 1.3617E-12 |

| | | | | | |
|---|---|---|---|---|---|
| CITED2 | 1.05076E-38 | 0.342704833 | 0.274 | 0.087 | 2.6022E-34 |
| BCLAF1 | 4.49032E-30 | 0.343373727 | 0.518 | 0.288 | 1.112E-25 |
| DUSP6 | 3.39952E-13 | 0.344148715 | 0.584 | 0.445 | 8.4189E-09 |
| AL596442.2 | 1.18152E-75 | 0.344250622 | 0.242 | 0.032 | 2.926E-71 |
| CREM | 1.43181E-25 | 0.344594164 | 0.412 | 0.213 | 3.5459E-21 |
| FCGRT | 4.40385E-20 | 0.345869578 | 0.783 | 0.701 | 1.0906E-15 |
| HSPE1 | 3.88378E-15 | 0.346285433 | 0.678 | 0.535 | 9.6182E-11 |
| DIS3 | 5.8068E-34 | 0.34721914 | 0.372 | 0.161 | 1.4381E-29 |
| SERTAD1 | 4.06908E-23 | 0.347257404 | 0.46 | 0.262 | 1.0077E-18 |
| ABCA9 | 2.09997E-43 | 0.347268274 | 0.34 | 0.111 | 5.2006E-39 |
| XBP1 | 1.33466E-33 | 0.347712094 | 0.405 | 0.181 | 3.3053E-29 |
| DDX21 | 1.71797E-23 | 0.349556604 | 0.564 | 0.356 | 4.2546E-19 |
| RAB20 | 5.4628E-52 | 0.350462535 | 0.409 | 0.138 | 1.3529E-47 |
| TOP1 | 2.06226E-27 | 0.351382141 | 0.549 | 0.324 | 5.1072E-23 |
| HMBOX1 | 8.42758E-38 | 0.353309219 | 0.391 | 0.165 | 2.0871E-33 |
| HS3ST3B1 | 8.4586E-55 | 0.354617186 | 0.295 | 0.074 | 2.0948E-50 |
| BSDC1 | 8.08075E-43 | 0.354995189 | 0.391 | 0.151 | 2.0012E-38 |
| RBM39 | 4.96841E-28 | 0.355618162 | 0.809 | 0.594 | 1.2304E-23 |
| AC015912.3 | 5.1428E-57 | 0.358330001 | 0.281 | 0.066 | 1.2736E-52 |
| SORBS1 | 6.48347E-52 | 0.359624724 | 0.384 | 0.127 | 1.6056E-47 |
| ZFAND5 | 3.8686E-20 | 0.359688111 | 0.656 | 0.445 | 9.5806E-16 |
| DAB2 | 8.65118E-26 | 0.36040636 | 0.626 | 0.405 | 2.1425E-21 |
| EGFL7 | 7.81092E-19 | 0.360582179 | 0.819 | 0.757 | 1.9344E-14 |
| VMO1 | 1.27728E-43 | 0.361952374 | 0.493 | 0.202 | 3.1632E-39 |
| H2AFX | 4.19267E-46 | 0.362002271 | 0.347 | 0.117 | 1.0383E-41 |
| KLF6 | 1.28405E-14 | 0.362721899 | 0.783 | 0.708 | 3.18E-10 |
| NOSTRIN | 6.22752E-30 | 0.362766559 | 0.595 | 0.353 | 1.5422E-25 |
| ABCG1 | 4.4204E-33 | 0.364909344 | 0.442 | 0.217 | 1.0947E-28 |
| BMPER | 1.55617E-49 | 0.365019798 | 0.437 | 0.16 | 3.8538E-45 |
| IFI16 | 3.70289E-20 | 0.365166984 | 0.709 | 0.531 | 9.1702E-16 |
| HSPA8 | 5.43127E-24 | 0.365721681 | 0.905 | 0.817 | 1.3451E-19 |
| PHLDA1 | 4.72312E-28 | 0.366222892 | 0.307 | 0.13 | 1.1697E-23 |
| KLF7 | 5.59338E-31 | 0.366762454 | 0.485 | 0.256 | 1.3852E-26 |
| POLR2J3.1 | 2.73372E-56 | 0.367121572 | 0.332 | 0.093 | 6.7701E-52 |
| IL18R1 | 1.33369E-69 | 0.368048891 | 0.243 | 0.036 | 3.3029E-65 |

| | | | | | |
|---|---|---|---|---|---|
| CYCS | 1.15911E-26 | 0.36806698 | 0.45 | 0.243 | 2.8705E-22 |
| MS4A4A | 1.1801E-46 | 0.368430506 | 0.353 | 0.115 | 2.9225E-42 |
| PLCG2 | 2.78752E-46 | 0.370141035 | 0.71 | 0.377 | 6.9033E-42 |
| RRBP1 | 3.54582E-27 | 0.370150356 | 0.612 | 0.392 | 8.7812E-23 |
| HSPD1 | 1.90866E-23 | 0.37032449 | 0.527 | 0.317 | 4.7268E-19 |
| PHC2 | 4.06814E-33 | 0.37146308 | 0.563 | 0.312 | 1.0075E-28 |
| MEF2C | 7.14155E-18 | 0.373302182 | 0.688 | 0.543 | 1.7686E-13 |
| PDE2A | 9.35788E-35 | 0.374122474 | 0.597 | 0.333 | 2.3175E-30 |
| MAN1C1 | 2.4566E-41 | 0.375274586 | 0.426 | 0.174 | 6.0838E-37 |
| GPATCH2L | 5.87382E-43 | 0.37545157 | 0.369 | 0.137 | 1.4547E-38 |
| KDM6B | 5.72756E-31 | 0.375845185 | 0.429 | 0.212 | 1.4184E-26 |
| SBNO2 | 2.77104E-44 | 0.376875275 | 0.456 | 0.191 | 6.8625E-40 |
| PLTP | 1.61597E-36 | 0.379575627 | 0.498 | 0.234 | 4.0019E-32 |
| UPP1 | 3.01512E-23 | 0.379672259 | 0.586 | 0.377 | 7.4669E-19 |
| CTTNBP2 | 4.01672E-72 | 0.380485118 | 0.299 | 0.057 | 9.9474E-68 |
| PCF11 | 4.38523E-37 | 0.383707115 | 0.406 | 0.18 | 1.086E-32 |
| DDX3Y | 3.96282E-38 | 0.38453516 | 0.388 | 0.159 | 9.8139E-34 |
| IFI44L | 4.04912E-26 | 0.385394675 | 0.518 | 0.301 | 1.0028E-21 |
| TNFAIP3 | 3.23457E-15 | 0.386533703 | 0.533 | 0.392 | 8.0104E-11 |
| HIPK2 | 1.06648E-31 | 0.386863644 | 0.592 | 0.342 | 2.6411E-27 |
| TBC1D4 | 3.65392E-43 | 0.387616363 | 0.488 | 0.211 | 9.0489E-39 |
| SMTN | 9.51363E-38 | 0.387761614 | 0.522 | 0.262 | 2.3561E-33 |
| GRASP | 2.66005E-19 | 0.387843985 | 0.611 | 0.448 | 6.5876E-15 |
| TPST2 | 1.3483E-39 | 0.389987416 | 0.496 | 0.238 | 3.3391E-35 |
| MT1X | 5.3707E-18 | 0.390364291 | 0.605 | 0.409 | 1.3301E-13 |
| FCHSD2 | 1.19497E-37 | 0.391822331 | 0.499 | 0.249 | 2.9593E-33 |
| TSN | 1.59948E-40 | 0.39370956 | 0.501 | 0.235 | 3.9611E-36 |
| ABLIM3 | 5.09493E-62 | 0.394627217 | 0.383 | 0.113 | 1.2618E-57 |
| FAM133B | 8.05084E-42 | 0.394634551 | 0.417 | 0.174 | 1.9938E-37 |
| SRSF4 | 2.58084E-33 | 0.394644815 | 0.524 | 0.291 | 6.3915E-29 |
| ACER3 | 1.19915E-57 | 0.395110451 | 0.367 | 0.11 | 2.9697E-53 |
| SECISBP2 | 4.21679E-36 | 0.395166542 | 0.381 | 0.164 | 1.0443E-31 |
| ARHGAP26 | 1.13772E-51 | 0.39525333 | 0.417 | 0.15 | 2.8176E-47 |
| ZNF207 | 5.76874E-31 | 0.395804383 | 0.516 | 0.296 | 1.4286E-26 |
| WIPI1 | 1.14268E-58 | 0.397904189 | 0.402 | 0.124 | 2.8299E-54 |

| | | | | | |
|---|---|---|---|---|---|
| APOH | 2.07121E-16 | 0.399675522 | 0.157 | 0.063 | 5.1293E-12 |
| SNAI1 | 2.3837E-43 | 0.399833988 | 0.398 | 0.156 | 5.9032E-39 |
| PDLIM3 | 7.43779E-47 | 0.400474861 | 0.316 | 0.098 | 1.842E-42 |
| DGKE | 9.59609E-37 | 0.400585169 | 0.411 | 0.181 | 2.3765E-32 |
| ZBTB7A | 1.08206E-31 | 0.401323075 | 0.426 | 0.214 | 2.6797E-27 |
| EHD3 | 1.13629E-44 | 0.401524444 | 0.507 | 0.215 | 2.814E-40 |
| CD274 | 1.66474E-71 | 0.401872609 | 0.333 | 0.072 | 4.1227E-67 |
| ENG | 4.00555E-19 | 0.401918023 | 0.792 | 0.749 | 9.9197E-15 |
| EZR | 2.55464E-31 | 0.403080318 | 0.38 | 0.17 | 6.3266E-27 |
| MERTK | 1.33946E-54 | 0.403803138 | 0.4 | 0.135 | 3.3172E-50 |
| BTNL9 | 9.02745E-48 | 0.404077081 | 0.516 | 0.22 | 2.2356E-43 |
| ITGA1 | 1.87364E-32 | 0.40457403 | 0.53 | 0.28 | 4.6401E-28 |
| F2R | 1.27808E-20 | 0.404940531 | 0.656 | 0.496 | 3.1652E-16 |
| APP | 2.51176E-18 | 0.406819886 | 0.783 | 0.768 | 6.2204E-14 |
| AMBP | 4.39606E-13 | 0.406864148 | 0.155 | 0.071 | 1.0887E-08 |
| SRSF2 | 2.08647E-31 | 0.406943209 | 0.614 | 0.371 | 5.1671E-27 |
| MCL1 | 1.02354E-34 | 0.407797953 | 0.862 | 0.634 | 2.5348E-30 |
| SLC20A1 | 1.23362E-38 | 0.407927803 | 0.417 | 0.179 | 3.0551E-34 |
| CDKN1A | 4.35097E-32 | 0.410891213 | 0.588 | 0.342 | 1.0775E-27 |
| SC5D | 1.61182E-44 | 0.414554153 | 0.465 | 0.192 | 3.9917E-40 |
| RNF115 | 5.53761E-38 | 0.416374894 | 0.515 | 0.259 | 1.3714E-33 |
| PRICKLE2 | 3.19334E-65 | 0.418694414 | 0.38 | 0.106 | 7.9083E-61 |
| CETP | 2.13259E-39 | 0.41976338 | 0.507 | 0.232 | 5.2814E-35 |
| ARID1B | 2.61918E-36 | 0.420220326 | 0.46 | 0.229 | 6.4864E-32 |
| WARS | 4.45856E-24 | 0.420603774 | 0.66 | 0.477 | 1.1042E-19 |
| HNRNPU | 8.22866E-27 | 0.423015637 | 0.778 | 0.572 | 2.0378E-22 |
| RASD1 | 1.69114E-48 | 0.423270101 | 0.329 | 0.101 | 4.1881E-44 |
| ANGPTL4 | 1.78724E-30 | 0.423272064 | 0.371 | 0.167 | 4.4261E-26 |
| SRSF6 | 2.98279E-46 | 0.426080113 | 0.434 | 0.179 | 7.3869E-42 |
| SEMA6A | 1.49165E-38 | 0.427714529 | 0.564 | 0.303 | 3.6941E-34 |
| RBMS1 | 8.67889E-26 | 0.427891762 | 0.571 | 0.374 | 2.1493E-21 |
| JUNB | 1.18683E-42 | 0.428066997 | 0.952 | 0.821 | 2.9392E-38 |
| GDF15 | 4.11717E-30 | 0.428257704 | 0.206 | 0.064 | 1.0196E-25 |
| BAIAP2 | 1.23087E-54 | 0.429285482 | 0.454 | 0.17 | 3.0483E-50 |
| LGALS3BP | 6.56122E-56 | 0.429293754 | 0.535 | 0.21 | 1.6249E-51 |

| | | | | | |
|---|---|---|---|---|---|
| PPP1R15A | 7.60122E-38 | 0.429854451 | 0.73 | 0.451 | 1.8824E-33 |
| HSP90AB1 | 9.55716E-27 | 0.430074711 | 0.94 | 0.91 | 2.3668E-22 |
| THBD | 1.61322E-19 | 0.430379006 | 0.532 | 0.353 | 3.9951E-15 |
| FKBP5 | 1.81368E-34 | 0.43043368 | 0.617 | 0.35 | 4.4916E-30 |
| ST6GAL1 | 3.50269E-37 | 0.431923136 | 0.642 | 0.363 | 8.6744E-33 |
| SAA1 | 8.45954E-18 | 0.432020335 | 0.112 | 0.033 | 2.095E-13 |
| WSB1 | 6.10305E-33 | 0.432488516 | 0.707 | 0.456 | 1.5114E-28 |
| HSPA5 | 2.81824E-27 | 0.433872982 | 0.679 | 0.46 | 6.9794E-23 |
| HEXIM1 | 1.13219E-48 | 0.434138315 | 0.394 | 0.143 | 2.8039E-44 |
| OSMR | 2.02482E-38 | 0.436187002 | 0.484 | 0.235 | 5.0145E-34 |
| RNF4 | 4.34927E-48 | 0.437342259 | 0.439 | 0.177 | 1.0771E-43 |
| TAL1 | 1.96002E-62 | 0.438264766 | 0.384 | 0.113 | 4.854E-58 |
| UBE2D3 | 1.52215E-31 | 0.440515648 | 0.724 | 0.499 | 3.7696E-27 |
| SRGAP1 | 4.11699E-73 | 0.441682965 | 0.394 | 0.105 | 1.0196E-68 |
| HERPUD1 | 1.31183E-41 | 0.442165552 | 0.577 | 0.295 | 3.2487E-37 |
| CLK1 | 4.17613E-42 | 0.445937919 | 0.54 | 0.264 | 1.0342E-37 |
| IDI1 | 5.48113E-42 | 0.446540111 | 0.535 | 0.263 | 1.3574E-37 |
| EPOR | 1.35702E-70 | 0.44674319 | 0.417 | 0.118 | 3.3607E-66 |
| NXPE3 | 9.97331E-56 | 0.446766973 | 0.44 | 0.156 | 2.4699E-51 |
| TSPAN6 | 1.08214E-48 | 0.448257119 | 0.447 | 0.178 | 2.6799E-44 |
| TRIB1 | 2.76792E-51 | 0.449177292 | 0.344 | 0.109 | 6.8548E-47 |
| NUDT16 | 2.14657E-34 | 0.449884236 | 0.526 | 0.281 | 5.316E-30 |
| PIK3R3 | 2.83989E-27 | 0.450189293 | 0.625 | 0.418 | 7.033E-23 |
| AFF4 | 1.80285E-37 | 0.450981379 | 0.598 | 0.349 | 4.4648E-33 |
| BTG1 | 5.2971E-09 | 0.451403397 | 0.741 | 0.731 | 0.00013118 |
| ZC3HAV1 | 1.32496E-40 | 0.451626844 | 0.406 | 0.172 | 3.2813E-36 |
| TRA2A | 2.75126E-47 | 0.452903874 | 0.493 | 0.215 | 6.8135E-43 |
| IL6ST | 1.47763E-17 | 0.45293081 | 0.809 | 0.771 | 3.6593E-13 |
| SRSF7 | 1.61369E-33 | 0.453076095 | 0.789 | 0.597 | 3.9963E-29 |
| PTPRB | 2.7417E-19 | 0.454353802 | 0.761 | 0.688 | 6.7898E-15 |
| LPAR6 | 8.98031E-32 | 0.455136595 | 0.575 | 0.348 | 2.224E-27 |
| NUFIP2 | 3.14072E-36 | 0.455485789 | 0.55 | 0.298 | 7.778E-32 |
| SLC40A1 | 5.05409E-36 | 0.455898582 | 0.595 | 0.338 | 1.2516E-31 |
| NCOA7 | 1.00631E-33 | 0.457185736 | 0.637 | 0.392 | 2.4921E-29 |
| ERRFI1 | 2.33517E-65 | 0.458097269 | 0.417 | 0.126 | 5.783E-61 |

| | | | | | |
|---|---|---|---|---|---|
| PPFIBP1 | 1.93033E-30 | 0.459822255 | 0.685 | 0.488 | 4.7805E-26 |
| AMD1 | 8.96452E-43 | 0.462278349 | 0.47 | 0.209 | 2.2201E-38 |
| RALGAPA2 | 2.74113E-51 | 0.467215222 | 0.538 | 0.242 | 6.7884E-47 |
| AFF1 | 2.04537E-28 | 0.467433865 | 0.558 | 0.351 | 5.0654E-24 |
| NABP1 | 2.56624E-67 | 0.469700914 | 0.302 | 0.065 | 6.3553E-63 |
| PLSCR1 | 4.11824E-35 | 0.471678939 | 0.65 | 0.413 | 1.0199E-30 |
| CFP | 2.9238E-53 | 0.473607451 | 0.515 | 0.206 | 7.2408E-49 |
| DDX5 | 4.84397E-42 | 0.47442257 | 0.989 | 0.93 | 1.1996E-37 |
| DENND4C | 4.49897E-59 | 0.47618965 | 0.502 | 0.196 | 1.1142E-54 |
| CEBPB | 4.83529E-42 | 0.476924863 | 0.553 | 0.274 | 1.1975E-37 |
| KLF9 | 5.05893E-26 | 0.47717248 | 0.695 | 0.499 | 1.2528E-21 |
| RASIP1 | 1.15567E-30 | 0.47729469 | 0.633 | 0.428 | 2.862E-26 |
| ANKRD44 | 8.79884E-89 | 0.480285242 | 0.34 | 0.06 | 2.179E-84 |
| IVNS1ABP | 6.59998E-51 | 0.482755562 | 0.51 | 0.217 | 1.6345E-46 |
| IFRD1 | 3.86557E-51 | 0.484839273 | 0.464 | 0.189 | 9.5731E-47 |
| THUMPD3-AS1 | 3.69715E-62 | 0.485074148 | 0.394 | 0.123 | 9.156E-58 |
| PRKCA | 6.1306E-96 | 0.486231405 | 0.347 | 0.058 | 1.5182E-91 |
| SNRK | 7.44984E-29 | 0.489427366 | 0.619 | 0.414 | 1.845E-24 |
| TLR4 | 1.18029E-47 | 0.490048976 | 0.561 | 0.267 | 2.923E-43 |
| CHORDC1 | 5.01545E-59 | 0.491577139 | 0.431 | 0.15 | 1.2421E-54 |
| ICAM1 | 2.78553E-50 | 0.492372055 | 0.453 | 0.18 | 6.8984E-46 |
| HIF1A | 1.6689E-24 | 0.497031234 | 0.691 | 0.539 | 4.133E-20 |
| TNFRSF10D | 1.70741E-62 | 0.498130966 | 0.46 | 0.161 | 4.2284E-58 |
| SOD2 | 2.69671E-36 | 0.498305049 | 0.482 | 0.244 | 6.6784E-32 |
| PLEKHG1 | 2.13597E-43 | 0.502639097 | 0.56 | 0.303 | 5.2897E-39 |
| F8 | 2.43797E-27 | 0.505725688 | 0.701 | 0.518 | 6.0376E-23 |
| NPL | 1.84918E-78 | 0.507862223 | 0.499 | 0.149 | 4.5795E-74 |
| LRG1 | 5.13255E-74 | 0.508101126 | 0.321 | 0.066 | 1.2711E-69 |
| SRGN | 4.19211E-42 | 0.510743407 | 0.938 | 0.846 | 1.0382E-37 |
| ZNF117 | 1.82896E-64 | 0.511963162 | 0.287 | 0.061 | 4.5294E-60 |
| APOC2 | 3.12331E-21 | 0.513929185 | 0.14 | 0.043 | 7.7349E-17 |
| GOLGB1 | 1.81976E-35 | 0.514358312 | 0.598 | 0.368 | 4.5066E-31 |
| EIF4A2 | 1.35929E-46 | 0.516782092 | 0.733 | 0.448 | 3.3663E-42 |
| DUSP1 | 4.7022E-46 | 0.516894548 | 0.963 | 0.857 | 1.1645E-41 |
| ITPRIP | 1.23202E-40 | 0.522863815 | 0.617 | 0.359 | 3.0511E-36 |

| | | | | | |
|---|---|---|---|---|---|
| NSUN6 | 9.546E-61 | 0.525300785 | 0.412 | 0.136 | 2.3641E-56 |
| RSRC2 | 9.79861E-52 | 0.526086176 | 0.577 | 0.281 | 2.4266E-47 |
| AKAP17A | 1.27451E-75 | 0.526599925 | 0.465 | 0.143 | 3.1563E-71 |
| HNRNPA2B1 | 5.80091E-47 | 0.527138753 | 0.963 | 0.845 | 1.4366E-42 |
| PRCP | 3.84529E-31 | 0.527373611 | 0.747 | 0.571 | 9.5229E-27 |
| UGCG | 1.05776E-35 | 0.529923313 | 0.56 | 0.324 | 2.6195E-31 |
| JUND | 8.39937E-43 | 0.530328799 | 0.978 | 0.91 | 2.0801E-38 |
| NOTCH4 | 4.86249E-45 | 0.531978474 | 0.693 | 0.417 | 1.2042E-40 |
| MALAT1 | 1.01911E-44 | 0.535988543 | 0.998 | 0.916 | 2.5238E-40 |
| NTN4 | 5.73245E-44 | 0.537025063 | 0.647 | 0.357 | 1.4196E-39 |
| DAAM1 | 4.3942E-47 | 0.539637044 | 0.538 | 0.261 | 1.0882E-42 |
| SRSF5 | 4.10208E-48 | 0.540019621 | 0.808 | 0.551 | 1.0159E-43 |
| ANKS1A | 1.86769E-59 | 0.540264268 | 0.476 | 0.18 | 4.6253E-55 |
| ANPEP | 3.70843E-70 | 0.543409143 | 0.487 | 0.162 | 9.1839E-66 |
| DDIT3 | 2.89425E-72 | 0.544053726 | 0.4 | 0.109 | 7.1676E-68 |
| EIF5 | 8.974E-44 | 0.544308686 | 0.664 | 0.39 | 2.2224E-39 |
| CUL4B | 1.20706E-38 | 0.545060524 | 0.428 | 0.2 | 2.9893E-34 |
| ZNF451 | 4.04884E-68 | 0.550637127 | 0.553 | 0.221 | 1.0027E-63 |
| IL1R1 | 1.23668E-45 | 0.552131988 | 0.684 | 0.392 | 3.0626E-41 |
| XAF1 | 1.41062E-56 | 0.553426385 | 0.636 | 0.31 | 3.4934E-52 |
| SFPQ | 1.54632E-49 | 0.555571165 | 0.746 | 0.49 | 3.8295E-45 |
| TSPYL2 | 1.90587E-73 | 0.558232622 | 0.364 | 0.09 | 4.7199E-69 |
| B4GALT1 | 9.66971E-45 | 0.559900202 | 0.622 | 0.358 | 2.3947E-40 |
| CXCL16 | 5.73712E-53 | 0.560547963 | 0.688 | 0.359 | 1.4208E-48 |
| GADD45B | 8.88731E-45 | 0.562800431 | 0.743 | 0.452 | 2.2009E-40 |
| ALDOB | 2.84254E-26 | 0.563314669 | 0.163 | 0.048 | 7.0395E-22 |
| PLAC8 | 1.61152E-52 | 0.563685209 | 0.507 | 0.206 | 3.9909E-48 |
| NPY1R | 2.64361E-65 | 0.564410526 | 0.544 | 0.211 | 6.5469E-61 |
| DDX3X | 9.67615E-47 | 0.565124827 | 0.848 | 0.597 | 2.3963E-42 |
| C1QTNF1 | 5.77963E-61 | 0.566100468 | 0.633 | 0.282 | 1.4313E-56 |
| ATF3 | 1.20185E-58 | 0.574371282 | 0.515 | 0.203 | 2.9764E-54 |
| C6orf62 | 7.99377E-45 | 0.576693104 | 0.536 | 0.274 | 1.9797E-40 |
| HNRNPH1 | 3.67945E-60 | 0.582701338 | 0.87 | 0.561 | 9.1122E-56 |
| GJA4 | 6.90544E-27 | 0.58369171 | 0.602 | 0.401 | 1.7101E-22 |
| LITAF | 3.24919E-50 | 0.587763138 | 0.617 | 0.313 | 8.0466E-46 |

| | | | | | |
|---|---|---|---|---|---|
| BRD2 | 8.02207E-49 | 0.588680807 | 0.614 | 0.323 | 1.9867E-44 |
| TCIM | 5.14482E-25 | 0.591671655 | 0.718 | 0.55 | 1.2741E-20 |
| TGFBR3 | 4.14362E-41 | 0.59278271 | 0.654 | 0.397 | 1.0262E-36 |
| IER2 | 2.83783E-48 | 0.594611607 | 0.926 | 0.719 | 7.0279E-44 |
| APOLD1 | 7.94726E-20 | 0.594787457 | 0.698 | 0.573 | 1.9681E-15 |
| YES1 | 1.19902E-34 | 0.59580739 | 0.65 | 0.439 | 2.9694E-30 |
| FZD4 | 2.94434E-41 | 0.599029033 | 0.665 | 0.418 | 7.2916E-37 |
| TFPI | 1.69501E-35 | 0.605630092 | 0.772 | 0.656 | 4.1977E-31 |
| SPRY4 | 2.23486E-62 | 0.607708789 | 0.566 | 0.247 | 5.5346E-58 |
| RBBP6 | 2.47222E-64 | 0.61219618 | 0.516 | 0.205 | 6.1225E-60 |
| VMP1 | 1.6906E-57 | 0.614702682 | 0.603 | 0.29 | 4.1868E-53 |
| FLT1 | 3.06938E-29 | 0.621339428 | 0.769 | 0.65 | 7.6013E-25 |
| TMEM37 | 5.75541E-83 | 0.639304157 | 0.642 | 0.235 | 1.4253E-78 |
| C5AR2 | 2.72282E-79 | 0.641379726 | 0.398 | 0.1 | 6.7431E-75 |
| RELN | 5.73337E-68 | 0.641433457 | 0.557 | 0.209 | 1.4199E-63 |
| HSPA1B | 1.77859E-43 | 0.644234665 | 0.811 | 0.576 | 4.4047E-39 |
| SELENOP | 3.73065E-52 | 0.650666367 | 0.882 | 0.655 | 9.239E-48 |
| KCNQ1OT1 | 5.23072E-60 | 0.650750458 | 0.391 | 0.125 | 1.2954E-55 |
| SLC23A2 | 1.4504E-106 | 0.655937971 | 0.437 | 0.092 | 3.592E-102 |
| DNAJA1 | 2.05853E-56 | 0.65823648 | 0.922 | 0.735 | 5.098E-52 |
| DNAJA4 | 5.31486E-91 | 0.662504331 | 0.391 | 0.086 | 1.3162E-86 |
| CCNL1 | 7.88987E-78 | 0.666302983 | 0.761 | 0.375 | 1.9539E-73 |
| ORM1 | 1.63655E-20 | 0.668720474 | 0.163 | 0.059 | 4.0529E-16 |
| SLCO4A1 | 1.97846E-88 | 0.673748228 | 0.464 | 0.124 | 4.8996E-84 |
| RSRP1 | 7.99614E-63 | 0.674316998 | 0.636 | 0.314 | 1.9802E-58 |
| CTSD | 5.69896E-52 | 0.677769913 | 0.816 | 0.591 | 1.4113E-47 |
| FOSB | 3.81565E-59 | 0.679954561 | 0.843 | 0.548 | 9.4495E-55 |
| STAB2 | 4.8405E-91 | 0.684876649 | 0.631 | 0.219 | 1.1987E-86 |
| CLEC1B | 2.81365E-46 | 0.684959905 | 0.73 | 0.401 | 6.968E-42 |
| TIMP1 | 9.6545E-30 | 0.693094841 | 0.803 | 0.775 | 2.3909E-25 |
| TACC1 | 2.82286E-32 | 0.696860444 | 0.738 | 0.601 | 6.9908E-28 |
| RBM3 | 1.92047E-46 | 0.698348881 | 0.657 | 0.42 | 4.756E-42 |
| SLC2A3 | 4.27021E-64 | 0.702658869 | 0.719 | 0.368 | 1.0575E-59 |
| NAMPT | 1.19878E-58 | 0.703339863 | 0.786 | 0.52 | 2.9688E-54 |
| ACP5 | 1.72158E-52 | 0.703354641 | 0.687 | 0.366 | 4.2635E-48 |

| | | | | | |
|---|---|---|---|---|---|
| CD4 | 6.57905E-59 | 0.705040461 | 0.674 | 0.344 | 1.6293E-54 |
| SRSF3 | 1.20897E-71 | 0.713935276 | 0.904 | 0.668 | 2.994E-67 |
| ENC1 | 4.25321E-81 | 0.716874687 | 0.454 | 0.129 | 1.0533E-76 |
| NR2F1 | 1.95634E-56 | 0.725799886 | 0.636 | 0.309 | 4.8449E-52 |
| LYVE1 | 3.68221E-51 | 0.727616253 | 0.555 | 0.253 | 9.119E-47 |
| KLF4 | 1.65216E-67 | 0.728194082 | 0.718 | 0.375 | 4.0916E-63 |
| ID2 | 2.05245E-70 | 0.730622047 | 0.628 | 0.272 | 5.0829E-66 |
| RBP4 | 1.1444E-19 | 0.732015492 | 0.16 | 0.059 | 2.8341E-15 |
| HSPH1 | 3.978E-69 | 0.732282963 | 0.577 | 0.246 | 9.8515E-65 |
| FGFR1 | 9.60092E-62 | 0.738209345 | 0.538 | 0.244 | 2.3777E-57 |
| SLC7A8 | 9.18E-105 | 0.738971896 | 0.561 | 0.155 | 2.273E-100 |
| IRAK3 | 1.14092E-78 | 0.739008489 | 0.53 | 0.196 | 2.8255E-74 |
| NFKBIZ | 5.12289E-90 | 0.740766714 | 0.577 | 0.198 | 1.2687E-85 |
| SMCHD1 | 1.3564E-62 | 0.746011495 | 0.637 | 0.32 | 3.3591E-58 |
| FGL2 | 2.38884E-44 | 0.746156149 | 0.592 | 0.319 | 5.916E-40 |
| PPP1R10 | 1.47403E-69 | 0.74707334 | 0.616 | 0.282 | 3.6504E-65 |
| MAF | 1.10585E-48 | 0.750711293 | 0.743 | 0.482 | 2.7386E-44 |
| HSPA1A | 7.95074E-51 | 0.753853502 | 0.862 | 0.725 | 1.969E-46 |
| ABHD5 | 2.02999E-91 | 0.759893914 | 0.49 | 0.144 | 5.0273E-87 |
| SH3BP5 | 7.86674E-48 | 0.762043489 | 0.695 | 0.465 | 1.9482E-43 |
| IL1RL1 | 1.04552E-50 | 0.762464743 | 0.22 | 0.044 | 2.5892E-46 |
| IL4R | 5.26161E-89 | 0.765695694 | 0.614 | 0.245 | 1.303E-84 |
| HYAL2 | 8.77908E-33 | 0.769643078 | 0.786 | 0.767 | 2.1741E-28 |
| CD36 | 2.41948E-48 | 0.775084603 | 0.747 | 0.476 | 5.9919E-44 |
| CEMIP2 | 3.52306E-82 | 0.778332481 | 0.721 | 0.344 | 8.7249E-78 |
| FOS | 4.01821E-76 | 0.782434547 | 0.961 | 0.755 | 9.9511E-72 |
| NID1 | 8.22184E-55 | 0.791871937 | 0.671 | 0.411 | 2.0361E-50 |
| RAPGEF5 | 8.13043E-47 | 0.793041493 | 0.713 | 0.525 | 2.0135E-42 |
| LIFR | 2.88756E-32 | 0.794789165 | 0.727 | 0.572 | 7.151E-28 |
| DLL1 | 4.35671E-86 | 0.797522276 | 0.569 | 0.205 | 1.0789E-81 |
| APOA1 | 1.28098E-09 | 0.817375727 | 0.203 | 0.127 | 3.1724E-05 |
| SLC38A2 | 4.23845E-62 | 0.817548223 | 0.806 | 0.585 | 1.0497E-57 |
| LGMN | 2.64269E-74 | 0.842979942 | 0.718 | 0.385 | 6.5446E-70 |
| FCN3 | 1.83469E-45 | 0.863280723 | 0.783 | 0.554 | 4.5436E-41 |
| PLIN2 | 4.04325E-82 | 0.869066617 | 0.718 | 0.351 | 1.0013E-77 |

| | | | | | |
|---|---|---|---|---|---|
| MRC1 | 2.99105E-54 | 0.873853366 | 0.741 | 0.493 | 7.4073E-50 |
| PLPP3 | 1.4226E-49 | 0.880842219 | 0.76 | 0.605 | 3.5231E-45 |
| ME2 | 2.97992E-93 | 0.889075954 | 0.56 | 0.187 | 7.3798E-89 |
| ZNF160 | 1.2053E-124 | 0.89741238 | 0.606 | 0.179 | 2.985E-120 |
| APOC3 | 9.75347E-14 | 0.898374181 | 0.216 | 0.118 | 2.4154E-09 |
| JUN | 3.04361E-90 | 0.913239412 | 0.963 | 0.773 | 7.5375E-86 |
| OIT3 | 5.62372E-68 | 0.937168379 | 0.73 | 0.395 | 1.3927E-63 |
| FCN2 | 7.11651E-63 | 0.943239121 | 0.752 | 0.419 | 1.7624E-58 |
| FCGR2B | 1.61369E-80 | 0.955754985 | 0.719 | 0.356 | 3.9963E-76 |
| TRA2B | 2.5045E-99 | 0.957254248 | 0.752 | 0.369 | 6.2025E-95 |
| AKAP12 | 2.72111E-56 | 0.958695052 | 0.761 | 0.503 | 6.7388E-52 |
| TTR | 3.43836E-17 | 0.959347508 | 0.188 | 0.085 | 8.5151E-13 |
| HSP90AA1 | 2.379E-107 | 0.97708194 | 0.98 | 0.918 | 5.892E-103 |
| MS4A6A | 4.23247E-85 | 0.989921464 | 0.735 | 0.357 | 1.0482E-80 |
| DUSP5 | 6.5147E-99 | 0.99218095 | 0.726 | 0.348 | 1.6134E-94 |
| CLEC4G | 3.77949E-72 | 1.008302343 | 0.752 | 0.406 | 9.3599E-68 |
| CRHBP | 1.41867E-59 | 1.018020686 | 0.753 | 0.484 | 3.5133E-55 |
| FUS | 1.2773E-106 | 1.019118386 | 0.784 | 0.391 | 3.163E-102 |
| DNASE1L3 | 3.59082E-58 | 1.022250526 | 0.786 | 0.584 | 8.8927E-54 |
| HP | 2.69677E-23 | 1.026542856 | 0.209 | 0.084 | 6.6786E-19 |
| DNAJB1 | 3.8036E-103 | 1.052503371 | 0.888 | 0.607 | 9.42E-99 |
| NEAT1 | 1.11028E-94 | 1.053949376 | 0.991 | 0.773 | 2.7496E-90 |
| MEG3 | 3.771E-99 | 1.066235437 | 0.459 | 0.112 | 9.3389E-95 |
| CD14 | 2.25934E-87 | 1.091555446 | 0.732 | 0.337 | 5.5953E-83 |
| SGK1 | 1.91255E-68 | 1.101761101 | 0.783 | 0.572 | 4.7364E-64 |
| CLEC4M | 5.31825E-91 | 1.124772627 | 0.682 | 0.283 | 1.3171E-86 |
| HES1 | 1.24727E-57 | 1.131742058 | 0.729 | 0.477 | 3.0889E-53 |
| PDK4 | 4.37748E-57 | 1.220373431 | 0.766 | 0.595 | 1.0841E-52 |
| STAB1 | 7.07659E-89 | 1.257936726 | 0.769 | 0.528 | 1.7525E-84 |
| **INTS6** | 1.1778E-134 | 1.366469603 | 0.674 | 0.232 | 2.917E-130 |
| EGR1 | 1.09108E-87 | 1.401191204 | 0.654 | 0.291 | 2.7021E-83 |
| ADM | 9.759E-109 | 1.412059404 | 0.726 | 0.338 | 2.417E-104 |
| CTSL | 1.63975E-77 | 1.412360562 | 0.786 | 0.568 | 4.0608E-73 |
| **ADAMTS4** | 1.24263E-95 | 1.498359432 | 0.679 | 0.319 | 3.0774E-91 |
| **MRO** | 6.4838E-110 | 1.754034026 | 0.657 | 0.25 | 1.606E-105 |
