## Supplementary material for "Multi-region spatial transcriptome analysis reveals cellular networks and pathways associated with hepatocellular carcinoma recurrence": Supp Fig

### Slide 1
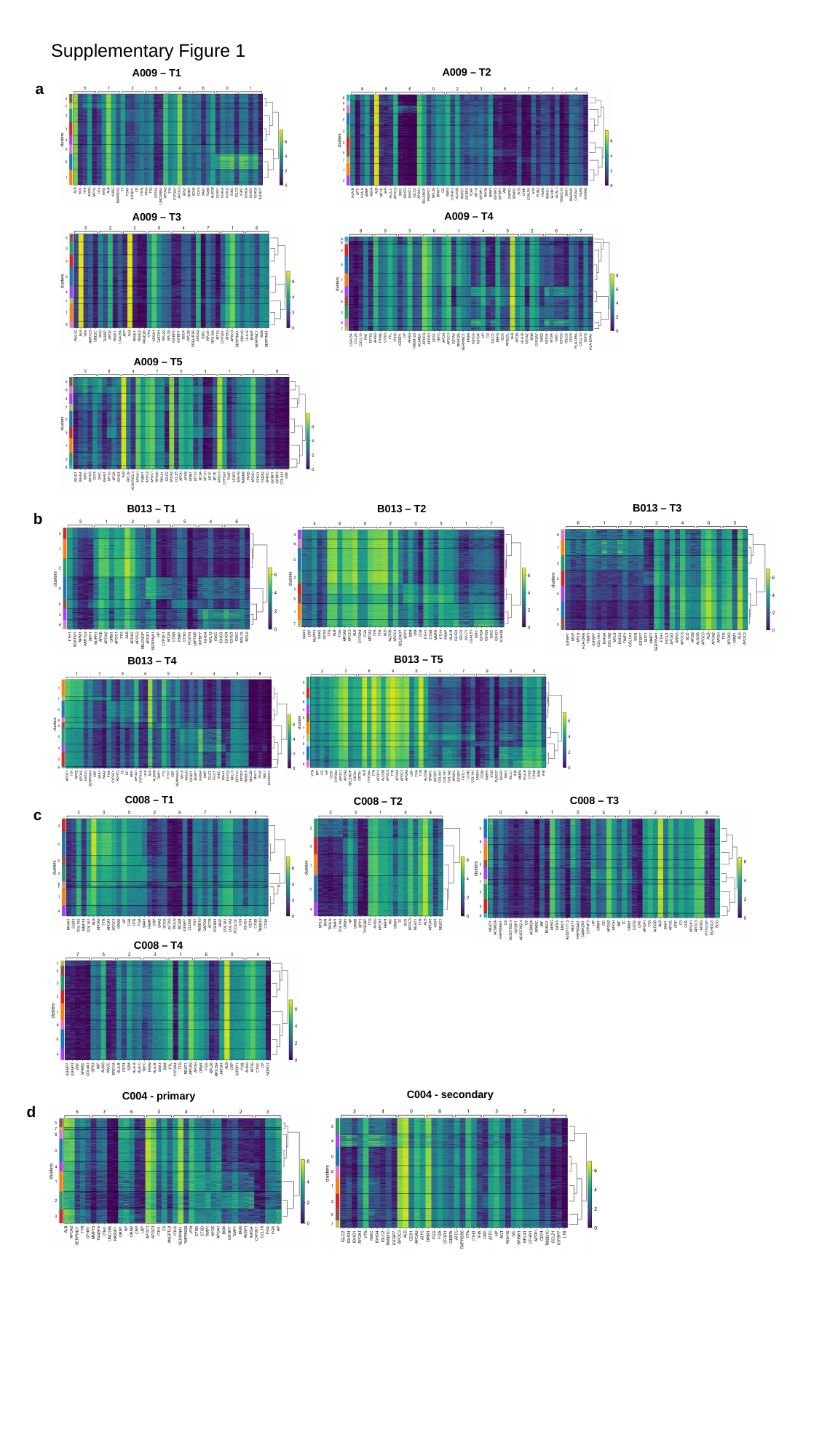

Supplementary Figure 1
A009 – T2
A009 – T1
a
A009 – T4
A009 – T3
A009 – T5
B013 – T3
B013 – T1
B013 – T2
b
B013 – T5
B013 – T4
C008 – T1
C008 – T3
C008 – T2
c
C008 – T4
C004 - secondary
C004 - primary
d

### Slide 2
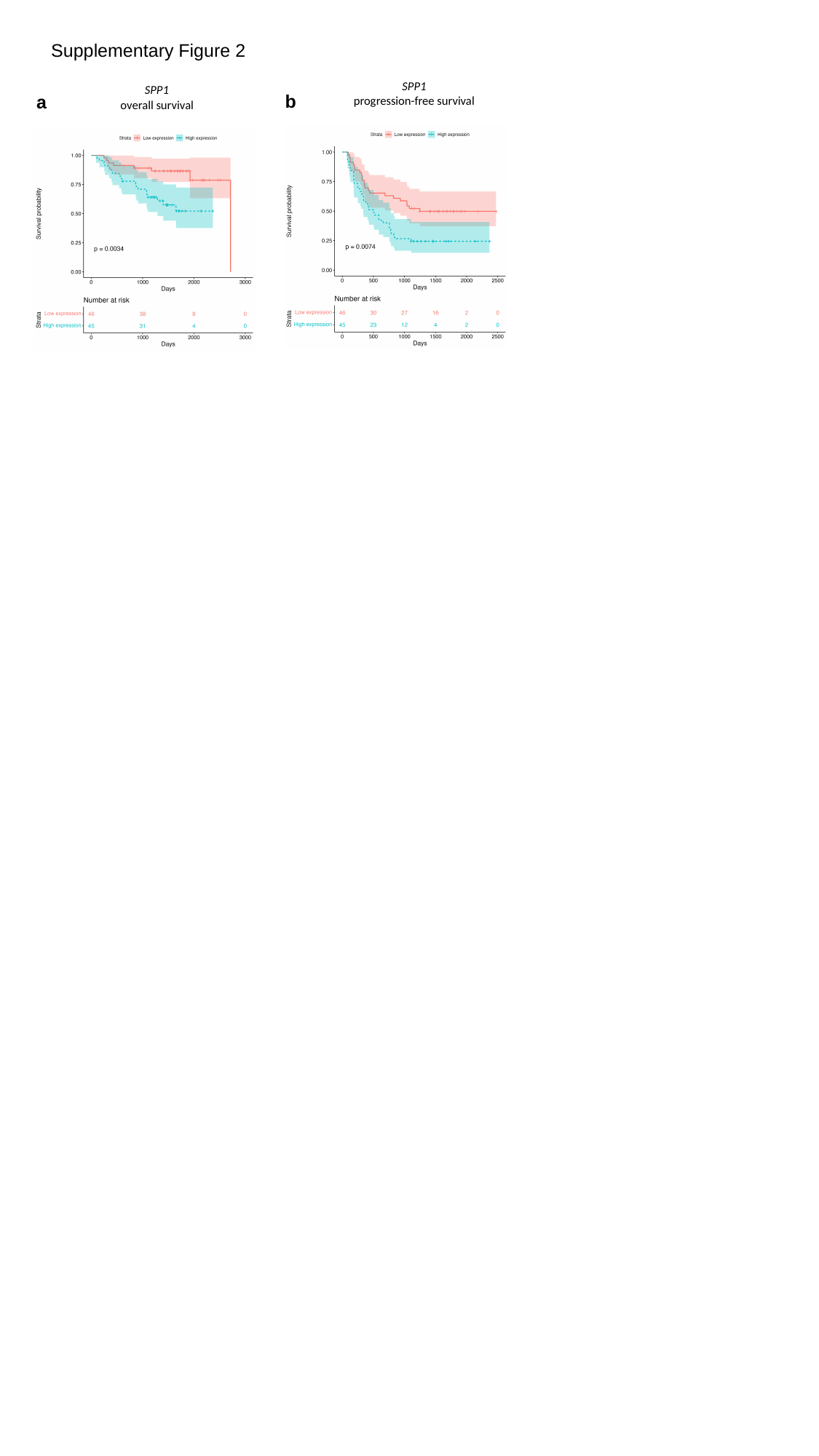

Supplementary Figure 2
SPP1
progression-free survival
SPP1
overall survival
b
a

### Slide 3
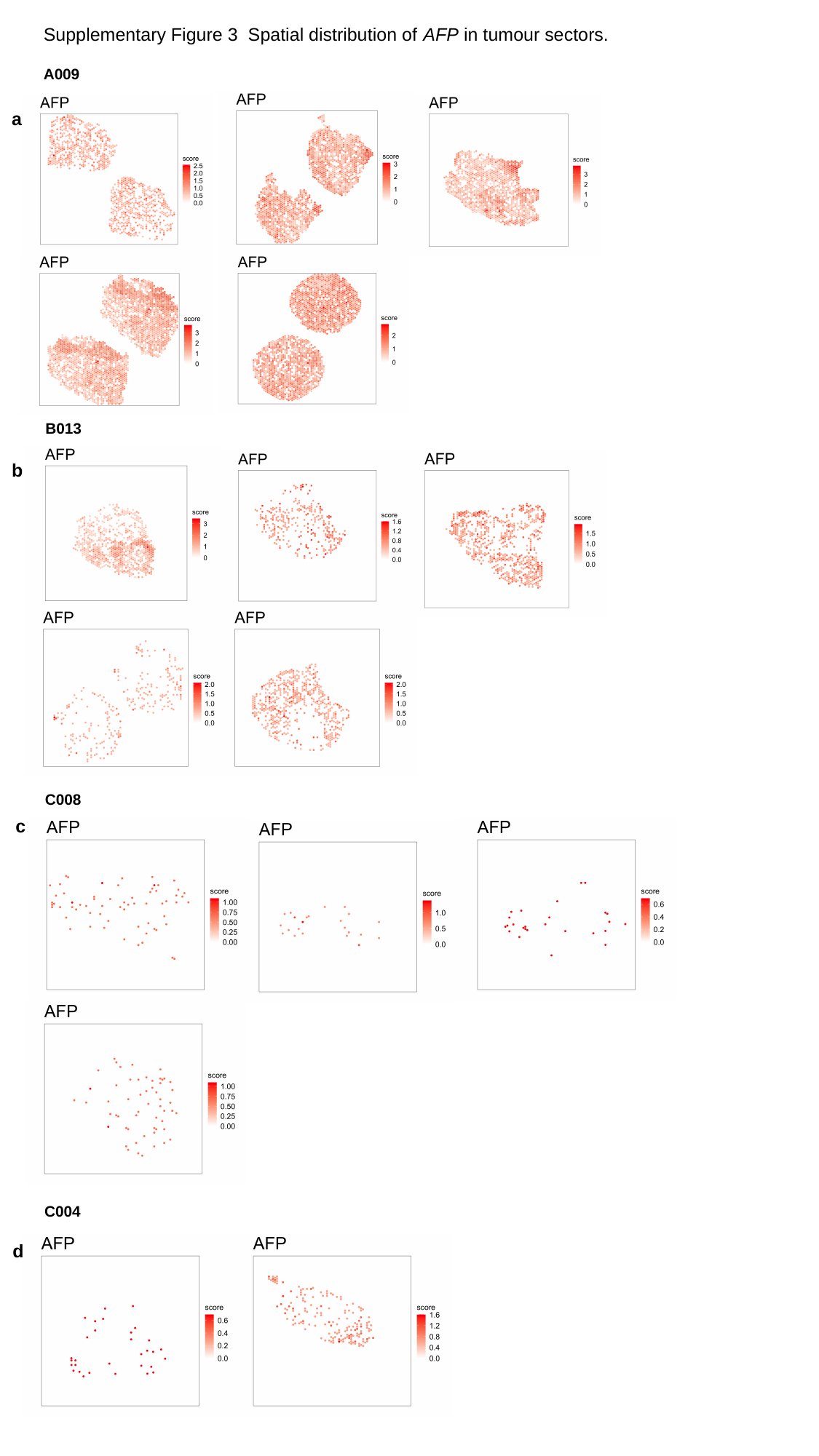

Supplementary Figure 3 Spatial distribution of AFP in tumour sectors.
A009
a
B013
b
C008
c
C004
d

### Slide 4
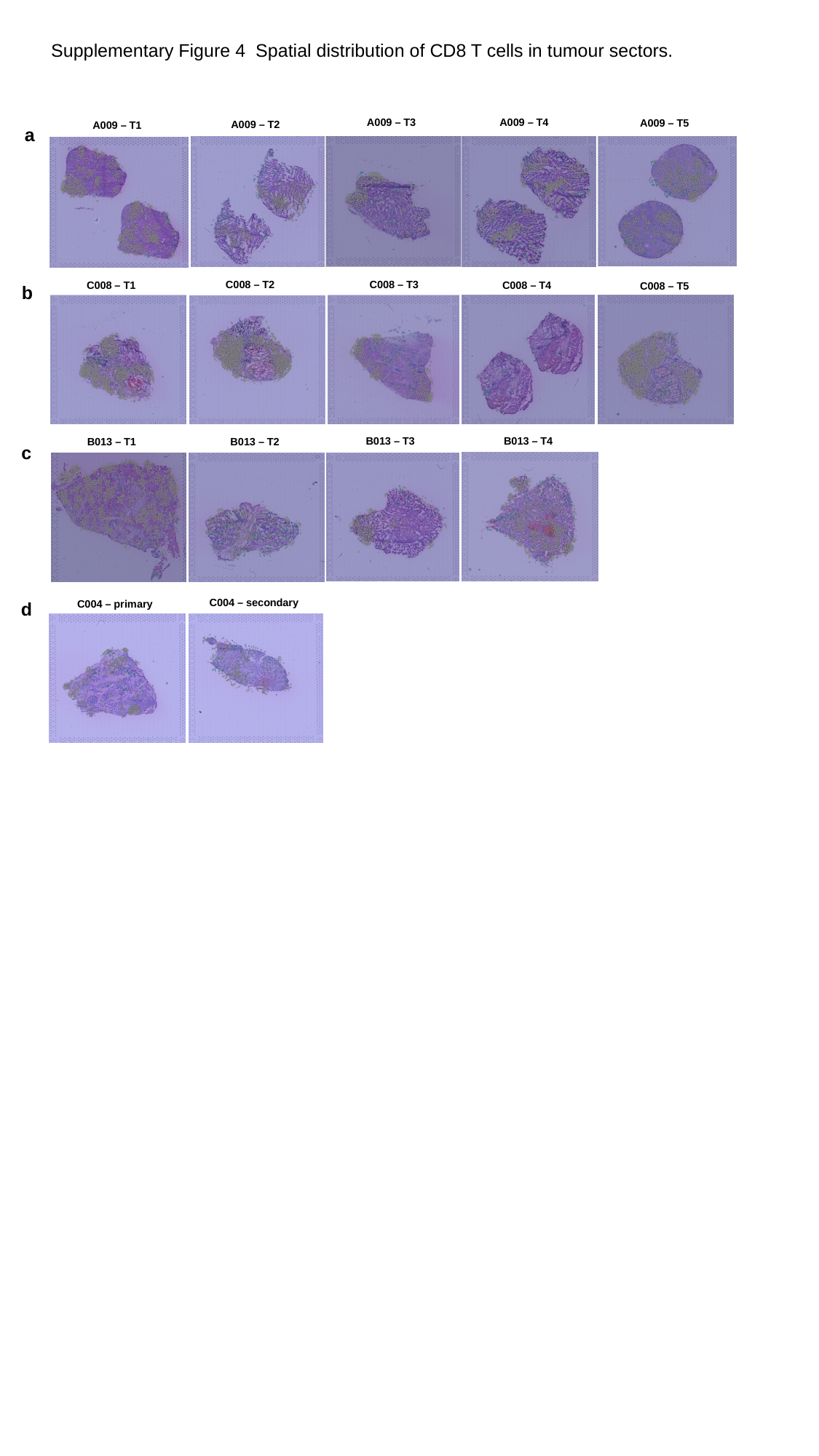

Supplementary Figure 4 Spatial distribution of CD8 T cells in tumour sectors.
A009 – T3
A009 – T4
A009 – T5
A009 – T2
A009 – T1
a
C008 – T3
C008 – T2
C008 – T4
C008 – T1
C008 – T5
b
B013 – T3
B013 – T4
B013 – T1
B013 – T2
c
C004 – secondary
C004 – primary
d
